# Diffusion tensor imaging of whole human brains during long-term formaldehyde fixation: Temporal evolution of diffusion parameters, post-mortem conditions, and dependence on tissue structure

**DOI:** 10.64898/2026.08.31.748254

**Authors:** Nina Lüthi, Francisco J. Fritz, Björn Fricke, Jan Malte Oeschger, Tobias Streubel, Herbert Mushumba, Klaus Püschel, Siawoosh Mohammadi

**Affiliations:** Center of Brain, Behaviour, and Metabolism, University of Luebeck, Luebeck, Germany; Department of Systems Neurosciences, University Medical Center Hamburg-Eppendorf, Hamburg, Germany; Department of Neuroradiology, University of Luebeck, Luebeck, Germany; Department of Neurophysics, Max Planck Institute for Human Cognitive and Brain Sciences, Leipzig, Germany; Department of Legal Medicine, University Medical Center Hamburg-Eppendorf, Hamburg, Germany

**Keywords:** diffusion MRI, DTI, immersion fixation, post-mortem time, ex vivo whole human brain

## Abstract

**Purpose:** Post-mortem Diffusion Magnetic Resonance Imaging (dMRI) findings on fixation-related changes in diffusion tensor imaging (DTI) parameters remain inconsistent, partly due to lower diffusion weighting and sparse early-fixation sampling. This study investigated how post-mortem tissue condition, fixation progression, and diffusion weighting shape DTI parameters in whole human brains, including whether an ex-vivo-adjusted protocol also suits in-situ and early fixation measurements.

**Methods:** Six neurologically healthy whole human brains (post-mortem interval (PMI) 12–24 h) were scanned across tissue conditions, including an in-vivo reference cohort. Five brains were scanned longitudinally through immersion fixation (0–150 days), yielding 175 datasets with dense early sampling. Diffusion measurements at b-values 1000–4000 s/mm^2^ assessed the influence of diffusion weighting on DTI parameters. Analyses covered multiple white- and gray-matter regions and within white matter stratification by fiber orientation dispersion (*κ*).

**Results:** The largest mean diffusivity (MD) shift occurred between in-vivo and in-situ, whereas fractional anisotropy (FA) changed most strongly during early fixation. During immersion fixation, MD showed reproducible monoexponential saturation, while FA was heterogeneous across brains. PMI, *κ*, and regional anatomy explained substantial FA heterogeneity across specimens. Qualitatively, the fixation-related changes matched across diffusion weightings, but precision across specimens was highest at b-value 4000 s/mm^2^.

**Conclusion:** During fixation, MD is a robust temporal marker, whereas FA requires microstructure-aware interpretation. Diffusion weighting modulates how clearly fixation effects can be resolved, but the same qualitative fixation dynamics remained detectable at in-vivo-like weighting with reduced precision.

## 1 Introduction

Conventional magnetic resonance (MR) images are acquired at millimeter-scale voxel resolution, whereas biologically relevant features of brain microstructure are organized at far smaller length scales. Diffusion Magnetic Resonance Imaging (dMRI) bridges this gap by probing restricted and direction-dependent water motion, with characteristic displacements of *∼* 1–10 *µ*m, two to three orders of magnitude below millimeter-size imaging voxels and is thereby sensitive to microstructural features that conventional MR contrast cannot resolve. [1, 2, 3] A widely used framework for analyzing diffusion-weighted images is diffusion tensor imaging (DTI), which uses the directional variation of water diffusion to estimate diffusion parameters that reflect local tissue organization. [4, 5] In the human brain, these parameters provide biologically and clinically relevant information about tissue composition and enable non-invasive in-vivo assessment of microstructure, with applications in neurological conditions such as neurodegenerative diseases [6, 7], epilepsy, [8] acute stroke, [9] and multiple sclerosis, [10] as well as psychiatric disorders [11, 12] and basic neuroscience. [13, 14]

To relate DTI parameters to underlying microstructure, ex-vivo histology is commonly used as the gold standard, enabling direct comparison of ex-vivo dMRI with quantitative histology and electron microscopy to validate sensitivity to axonal-signal fraction [15, 16] and caliber variation. [17, 3] Relations established ex-vivo may then be translated to in-vivo dMRI, [18] but cautiously, because physical and biochemical brain-tissue properties change substantially between in-vivo and ex-vivo conditions. [19]

After death, brain tissue undergoes biological alterations that progressively affect microstructural integrity, termed autolysis. Chemical fixation interrupts this degradation and preserves post-mortem samples. The post-mortem interval (PMI), defined as the interval between death and the start of fixation, is critical because autolytic changes during this period may affect ex-vivo dMRI measurements. [19] In addition, tissue is no longer in the living physiological state: loss of blood flow and oxygen supply, together with lower temperature, reduce water mobility, so diffusivity is typically lower ex-vivo than in-vivo. [20] This lower ex-vivo diffusivity, typically 2- to 5-fold below in-vivo levels, [19] requires protocol adaptation, primarily through the b-value. In-vivo DTI is commonly acquired at b-value 1000 s/mm^2^, [21, 22] reflecting a practical balance between contrast, scan time, and signal-to-noise ratio (SNR), whereas the ISMRM Diffusion Study Group recommends adapting ex-vivo b-values so that the product of b-value and tissue diffusivity is approximately one, placing ex-vivo DTI in the range 2000–5000 s/mm^2^. [19, 23] Higher diffusion weighting can restore comparable attenuation in ex-vivo tissue, but also reduces SNR, so protocol choice must be considered together with post-mortem tissue state rather than in isolation.

Immersion fixation adds further uncertainty because fixative enters from the tissue surface and penetrates large samples slowly. It is therefore unclear when a whole human brain can be considered fully fixed. Reported durations range from approximately 2 weeks to 30 days or longer, while early fixation time points remain sparsely sampled. [19, 24, 25, 26] Because penetration starts at tissue boundaries, spatial differences in MR parameter changes during fixation have often been interpreted as depth dependence, with deeper regions assumed to fix later or less completely than superficial regions. [27, 24] However, distance from the surface covaries with tissue composition and fiber architecture, and it remains unclear how much fixation-related DTI changes reflect geometric depth versus microstructure. Fiber orientation dispersion (*κ*) may therefore help explain fixation-induced DTI changes independently of geometric depth.

Uncertainty in fixation timing is further complicated by the diffusion protocol used to measure it. Previous work reported no fixation influence on diffusion parameters with in-vivo-like diffusion weighting, [28] whereas Dyrby et al. [29] proposed an ex-vivo-adjusted b-value 4000 s/mm^2^ for fixed porcine brains, optimized for fiber-orientation reconstruction rather than temporal-evolution precision of DTI parameters. [29, 30] Furthermore, their study did not target densely sampled early immersion-fixation kinetics. It therefore remains unclear whether the recommended ex-vivo b-value range provides sufficient sensitivity and precision for tracking DTI temporal evolution during fixation in whole human brains.

This study quantifies how post-mortem tissue state, fixation progression, and diffusion weighting shape the detectability, precision, and interpretability of DTI-derived FA and MD in whole human brains. Instead of treating ex-vivo tissue as a single condition, we distinguish post-mortem conditions spanning in-situ (the brain after death but still within the skull before extraction), fixation onset and progression, and subsequent hydration, covering both PMI-related changes and chemical fixation. We do not aim to define a universally applicable b-value, but test whether pragmatic diffusion-weighting choices can render fixation-related changes effectively unresolved in DTI parameters. We first compare FA and MD across these conditions. Second, we assess whether fixation-related DTI changes across b-values from 1000 to 4000 s/mm^2^ are detectable, and how reliably fixation trajectories can be resolved and modeled across diffusion weightings. Third, we examine whether PMI, regional anatomy, and *κ* explain inter-brain and inter-region heterogeneity in FA and MD during fixation beyond tissue depth alone. To address these aims, we acquired longitudinal dMRI in six neurologically healthy whole-brain donors from in-situ through densely sampled fixation and hydration, complemented by an in-vivo reference cohort.

## 2 Methods

### 2.1 Specimens and Sample preparation

#### In-vivo cohort

Four healthy participants (three male, one female; 35.75 *±* 9 years) were screened for neurological or psychiatric illness and studied with approval of the local ethics committee (Ärztekammer Hamburg, #PV51141) in accordance with the Declaration of Helsinki (seventh revision, 2013).

#### Ex-vivo specimens

Six whole, neurologically healthy human brains (four male, two female, 60.0 *±* 11 years) from deceased donors were used with prior informed consent (WF-74/16). PMIs ranged from 12 to 24 h; until autopsy the bodies were kept at a stable 18 °C to reduce tissue degeneration. [31] Of the six brains, five were scanned in situ before autopsy and five had longitudinal formaldehyde-fixation scans (one specimen lacked in-situ data, another lacked longitudinal fixation data); all six contributed to tissue-condition analyses where scans were available. Brains were immersion-fixed in 4 % formaldehyde, stored at 7 °C, and later rehydrated in PBS (solution compositions in Supporting Information, Section 7.1; scan timing per brain in Appendix Table A1). For thermal stability, brains were kept at room temperature for some hours before scanning, with 30 min breaks after each diffusion measurement. [32, 33, 34]

### 2.2 Image Acquisition

All measurements were performed on a 3 T PRISMA fit MRI (Siemens Healthcare, Erlangen, Germany). Ex-vivo acquisitions used a Siemens 32-channel receiver head coil and in-vivo a Siemens 64-channel coil. Diffusion data were acquired with an echo-planar pulse-gradient spin echo sequence.

#### Ex-vivo diffusion MRI

For each ex-vivo measurement, two multishell acquisitions were performed: 1k2k (b-shells 0, 1000, and 2000 s/mm^2^) and 2k4k (b-shells 0, 2000, and 4000 s/mm^2^), providing lower and higher ex-vivo diffusion weighting. The 2k4k acquisition covered the range recommended for formaldehyde-fixed tissue (approximately 2000–5000 s/mm^2^). [19] Acquisition parameters are provided in Table 1.

**Table 1:** Acquisition parameters and analyzed b-shells for the two ex-vivo multishell acquisitions (1k2k and 2k4k) and the in-vivo reference. Sequence parameters, including field of view (FOV) and GRAPPA 2, were constant between the ex-vivo acquisitions, whereas TE and TR differed. The 2k4k acquisition included blip-up and blip-down images; 1k2k included blip-up images only.

|  | 1k2k acquisition | 2k4k acquisition | In-vivo acquisition |
| --- | --- | --- | --- |
| repetition time (TR) | brain 1, 3-5: 5200 ms | 7000 ms | 3100 ms |
| echo time (TE) | 71 ms | 79 ms | 74.6 ms |
| slices | 96 | 96 |  |
| resolution<br>(isotropic voxel) | brain 1: (1.6 mm)<sup>3</sup><br>brain 3: (1.6 mm)<sup>3</sup><br>brain 4: (1.6 mm)<sup>3</sup><br>brain 5: (1.6 mm)<sup>3</sup><br>brain 6: (1.6 mm)<sup>3</sup> | brain 1: (1.6 mm)<sup>3</sup><br>brain 2: (2.5 mm)<sup>3</sup><br>brain 3: (1.6 mm)<sup>3</sup><br>brain 4: (1.6 mm)<sup>3</sup><br>brain 5: (1.6 mm)<sup>3</sup><br>brain 6: (1.6 mm)<sup>3</sup> | (1.7 mm)<sup>3</sup> |
| Acquired b-shells<br>[s/mm<sup>2</sup>]<br>(# directions) | 0 (15)<br>1000±50 (60)<br>2000±50 (60) | 0 (15)<br>2000±50 (60)<br>4000±50 (60) | 0 (11)<br>1000 (29)<br>2000 (64) |
| Analyzed b-shells<br>[s/mm<sup>2</sup>] | 1000 (main text)<br>2000 (Supporting Information) | 2000 (main text)<br>4000 (main text) | 1000 (in-vivo reference) |

#### In-vivo diffusion MRI

In-vivo data were acquired on the same system with a multishell protocol adapted for in-vivo diffusivity (b-shells 0, 1000, and 2000 s/mm^2^, Table 1).

### 2.3 Image Processing

Each ex-vivo multishell acquisition was split by b-shell into single-shell datasets. Acquired and analyzed shells are listed in Table 1. DTI was fitted with the ACID toolbox [35] in SPM 12 [36] (Matlab R2021b [37]).

Pre-processing comprised eddy current and motion correction (ECMOCO) and susceptibility-distortion correction (hyperelastic susceptibility artifact correction (HySCO)). As the 1k2k acquisition lacked blip-down images, the mean b0 images (blip-up and blip-down) from 2k4k after ECMOCO served as reference for its susceptibility correction, and the resulting unwarped 1k2k blip-up image was used for the DTI fit. For 2k4k, unwarped blip-up and blip-down images were combined into a weighted-average input image. All datasets were fitted with Non-linear Least Squares (NLLS) without rician bias correction. Tensor-fit quality was summarized per voxel by the root-mean-square residual between measured and predicted diffusion-weighted signals (Supporting Information, Section 7.4).

Image registration was performed brain-wise, using the first acquisition as reference (in-situ for all brains except brain 5). multi-parametric mapping (MPM) acquisition, MTsat estimation, and the detailed coregistration pipeline are described in Fritz et al.[38]. MTsat maps, which offer higher resolution and contrast, were registered first, and the resulting transformation was applied to the corresponding diffusion data. The coregistration pipeline is illustrated in the Supporting Information (Figure S1).

### 2.4 Regions of interest (ROIs)

After longitudinal coregistration, each brain was segmented into white matter (WM), deep grey matter (dGM), and cortical grey matter (cGM), and 11 region of interests (ROIs) were identified within these classes. Brain-specific mean FA and MD maps of the coregistered images were used for segmentation. Tissue-class masks were obtained from SPM segmentation in MNI space (MNI) using tissue probability maps (TPMs) at a 95% probability threshold [36]. Inverse deformation fields projected atlas labels (JHU for white-matter tracts, Harvard-Oxford for gray matter) into specimen space, refined by FA/MD-guided selection on the mean maps and visual quality control.

In the main manuscript, analyses of protocol effects or variability were restricted to the corpus callosum (CC), whose high anatomical alignment and comparatively homogeneous fiber architecture reduce confounding by structural heterogeneity. [39] Analyses comparing anatomical regions additionally included the other ROIs.

### 2.5 Analysis

Analyses comprised discrete tissue condition comparisons (Section 2.5.1) and continuous fixation time-course modeling (Sections 2.5.2, 2.5.4, and 2.5.5). Tissue condition analyses used all six ex-vivo brains where stage-specific data were available; continuous fixation models used the five brains with longitudinal formaldehyde-fixation scans. For each brain, scan day, and region, the regional median FA or MD (across ROI voxels) was computed and plotted against fixation time.

#### 2.5.1 Diffusion parameters across tissue conditions

Post-mortem measurements were organized into four tissue conditions that separate early physiological change after death from later chemical fixation and rehydration. This staging underpins the comparison in Section 3.1, the ex-vivo protocol comparison across tissue conditions (Section 2.5.3 and Supporting Information, Section 7.4), and the discrete-stage PMI analysis (Section 2.5.5**(ii)**).

*—* **In-vivo (reference):** diffusion MRI from four healthy volunteers on the same scanner (Table 1). This cohort provides a population reference but is not matched to individual ex-vivo donors.

*—* **In-situ:** ex-vivo scan in the skull after death, performed immediately before brain extraction and immersion fixation, acquired near the end of the PMI interval (12–24 h).

*—* **Beginning of fixation:** whole-brain immersion in 4 % formaldehyde within 0–3 days after autopsy and fixation onset.

*—* **Fixed:** continued immersion in 4 % formaldehyde at 40–45 days.

*—* **Hydrated:** rehydration in PBS after transfer from formaldehyde (timing per brain in Appendix Table A1).

For each stage, voxelwise FA and MD were extracted in the CC and pooled across ex-vivo donors with available scans (Supporting Information Table S2). Stage boxplots show voxel-value distributions with the in-vivo median as reference and annotated relative changes.

#### 2.5.2 Temporal evolution fixation models and model selection

Fixation time courses of *y ∈ {*MD, FA*}* were fitted with three saturation models, expressed as special cases of a single biexponential form:

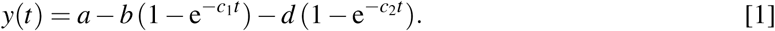

The biexponential model uses the full expression, with a fast and a slow compartmental process [40]. The monoexponential model sets *d* = 0, describing single-phase saturation consistent with formaldehyde cross-linking kinetics [25]. The null model sets *b* = *d* = 0, giving *y* = *a* (no temporal change).

The parameters have a direct physical meaning: *a* is the value at fixation onset (*y*(*t*_0_)); amplitudes *b* and *d* quantify the fast and slow components (positive values indicate a decrease); and *c*_1_*, c*_2_ are rate constants with time constants *τ^y^_i_* = 1*/c_i_* (days). After one *τ*, 63 % of that component’s amplitude is reached, so a larger *τ* indicates slower saturation. The model-amplitude *b* is distinct from the diffusion-weighting *b*-value.

Model selection used the corrected Akaike Information Criterion (AICc), evidence ratios, and a parsimony rule for similarly supported models (Supporting Information, Section 7.5.1). Fit quality was quantified by normalised root mean square error (nRMSE): for each b-value and parameter, brain–time medians were pooled and a robust amplitude Δ*y*^(*q*)^ = *P*_97.5_(*y*) *− P*_2.5_(*y*) was used, where *P*α (*y*) is the α-th percentile. For each model *m*,

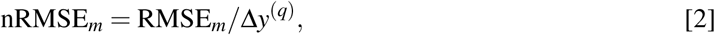

yielding a dimensionless residual comparable across parameters and protocols (Figure 3**G,H**).

**Figure 1:**
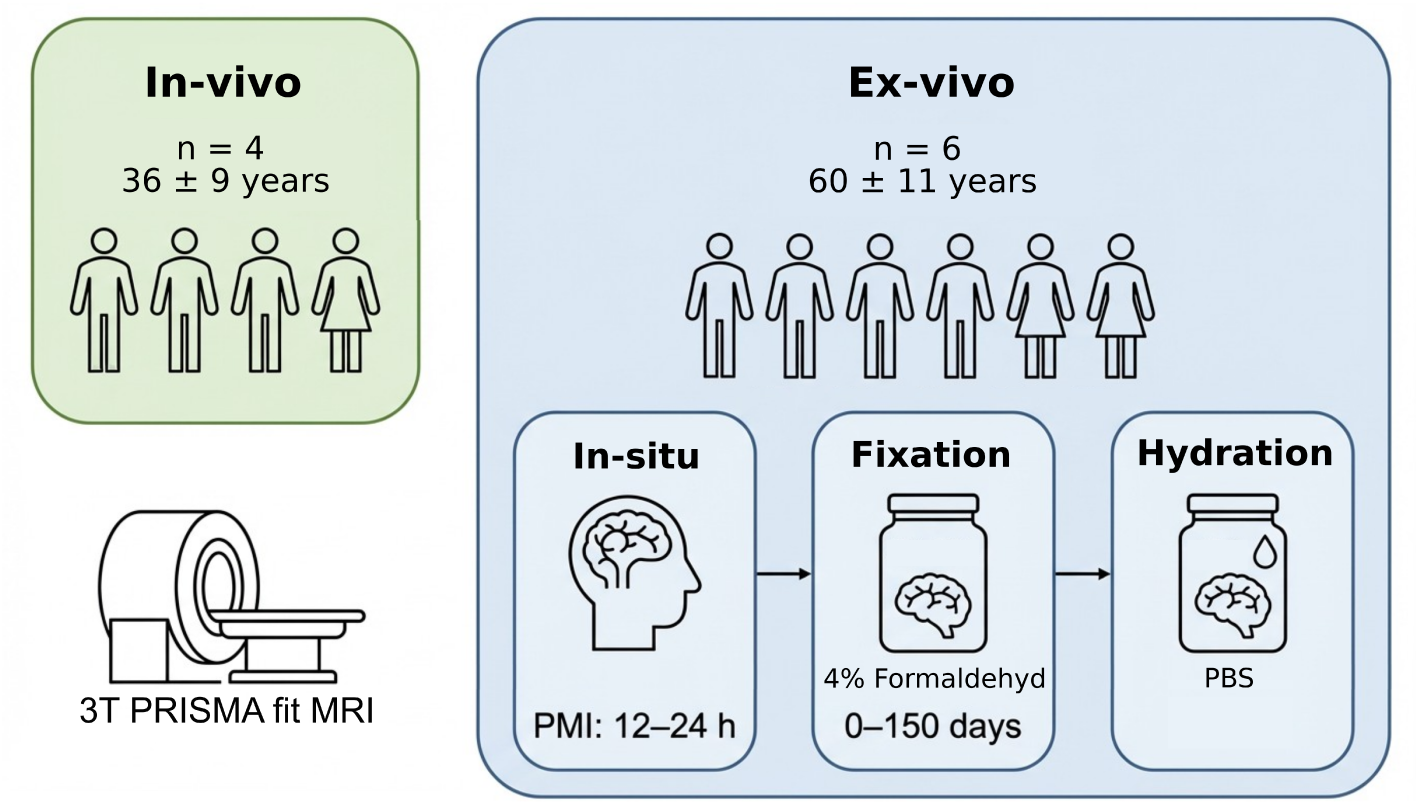
Study cohort and post-mortem acquisition workflow. In-vivo diffusion MRI was acquired in a reference cohort of four healthy volunteers (35.75 *±* 9 years) on a 3T Prisma fit system. Ex-vivo diffusion magnetic resonance imaging (MRI) was acquired in six whole human brain donors (60 *±* 11 years, post-mortem interval (PMI): 12–24 h) across successive post-mortem tissue states: in-situ (scanned after death but before brain extraction), immersion fixation in 4 % formaldehyde, and subsequent rehydration in phosphate-buffered saline (PBS). Five brains contributed longitudinal formaldehyde fixation data over 0–150 days. This design enables comparison between in-vivo reference measurements and the temporal evolution of diffusion parameters across post-mortem tissue conditions.

**Figure 2:**
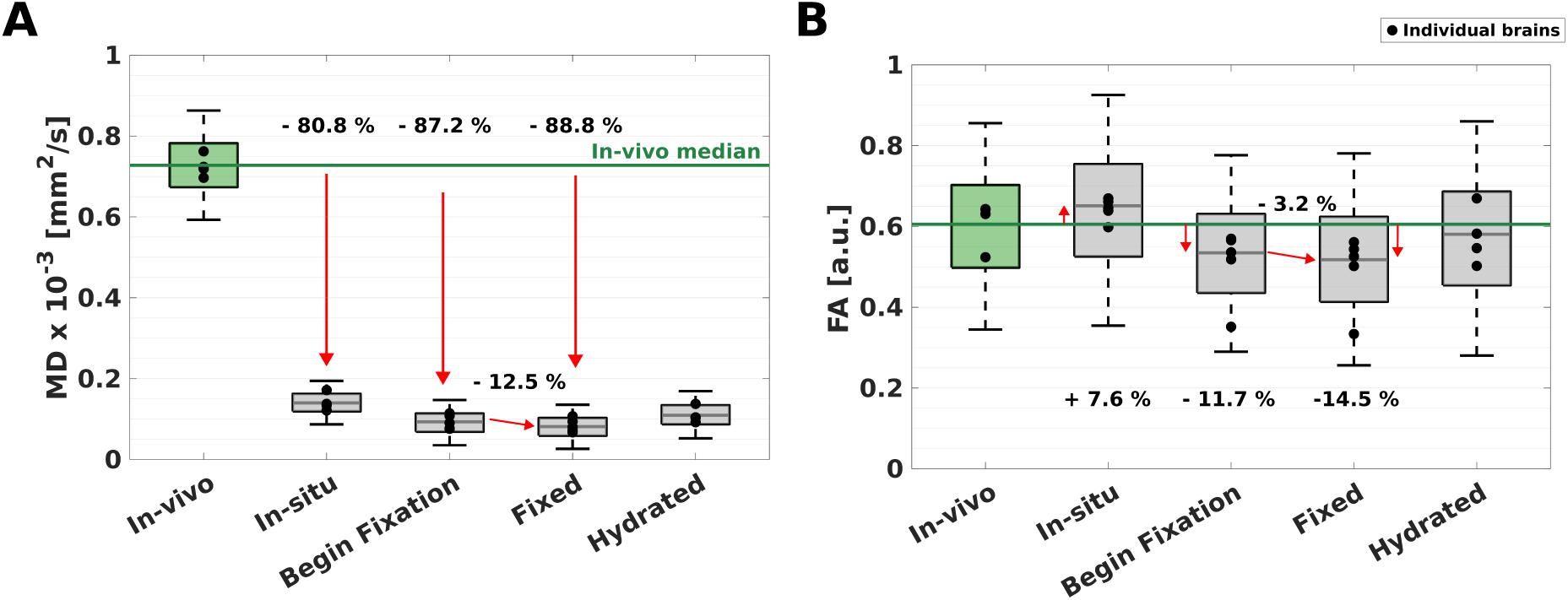
Diffusion parameters across tissue conditions in the corpus callosum (CC): Voxelwise mean diffusivity (MD) and fractional anisotropy (FA) in the corpus callosum (CC) are shown for five tissue conditions in (**A**) and (**B**), respectively: in-vivo (green), in-situ, beginning of fixation (0–3 days in 4 % formaldehyde), fixed (40–45 days in 4 % formaldehyde), and hydrated in phosphate-buffered saline (PBS). Boxplots show the voxelwise distribution pooled across all brains within each condition. Whiskers extend to the lowest and highest values within 0.75 times the interquartile range (IQR) (*±* 1.01 sigma, 68.76 % coverage of the data). The lower quartile (q_n_(0.25)) and upper quartile (q_n_(0.75)) bound the box (middle 50 % of values) and the center line indicates the median. The green box represents in-vivo data from four healthy subjects and gray boxes represent ex-vivo data from five whole human brains. A horizontal green line marks the in-vivo median as a reference. Individual brain medians are overlaid as markers. Annotated relative changes compare selected tissue condition transitions and selected ex-vivo conditions to the in-vivo median.

**Figure 3:**
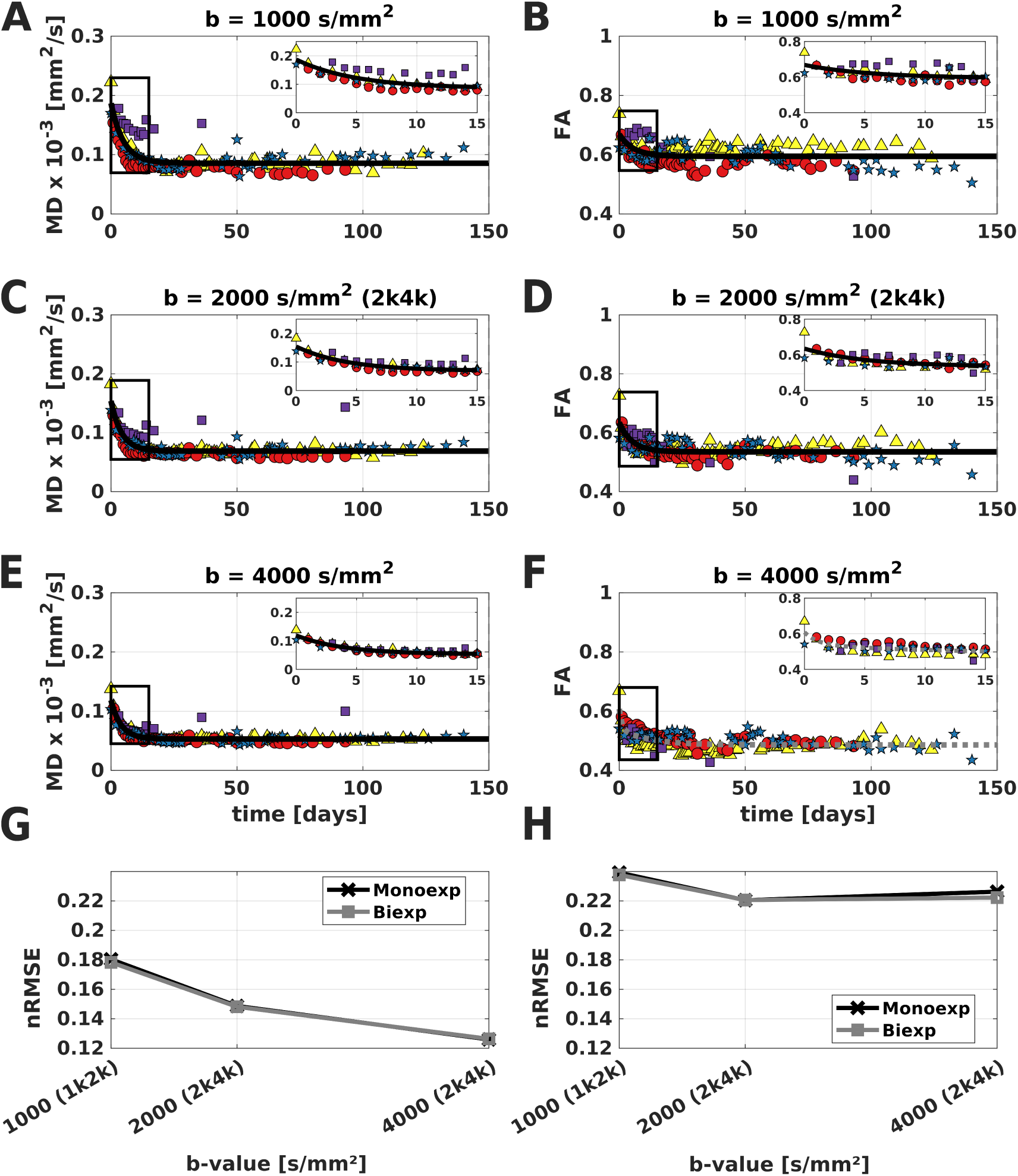
Temporal evolution modeling of mean diffusivity (MD) and fractional anisotropy (FA) during fixation: b-value comparison and model fit quality in the CC. Median MD (left column: **A**, **C**, **E**) and FA (right column: **B**, **D**, **F**) are shown over fixation time for *b* = 1000, 2000, and 4000 s/mm^2^, respectively. Each marker represents one brain-specific CC median at one fixation time point; color and marker shape indicate the specimen and post-mortem interval (PMI): brain3 (15 h, yellow triangle), brain6 (18 h, red circle), brain1 (21 h, purple square), and brain5 (24 h, blue star). Insets enlarge the early fixation phase (first 15 days). Overlaid curves show the preferred saturation model for each parameter and b-value, selected by AICc and evidence ratios (Supporting Information Table S3); monoexponential and biexponential alternatives are shown in Supporting Information 7.5. Panels **G** and **H** summarise normalised root mean square error (nRMSE) for MD and FA, respectively, across b-values; nRMSE is defined in Eq. 2.

#### 2.5.3 Protocol comparison and selection for main analyses

In the CC, three diffusion weightings were compared (b-value 1000 s/mm^2^ from 1k2k and b-values 2000 and 4000 s/mm^2^ from 2k4k, Table 1) for fixation-related changes, parameter dispersion, and model fit quality. Complementary signal-level diagnostics and the four-dataset comparison, including the additional b-value 2000 s/mm^2^ shell from 1k2k, are in Supporting Information (Sections 7.4.1, 7.4.1, and 7.4.2). Subsequent main analyses used b-value 4000 s/mm^2^, which had the lowest parameter variance across tissue conditions.

For each brain and b-value, median FA and MD per fixation time point (over CC voxels) were fitted with the saturation models in Section 2.5.2. WM results (median over WM voxels per time point) are in the Supporting Information (Figures S7 and S6).

#### 2.5.4 Temporal evolution fixation dynamics in the corpus callosum

Only time points during immersion in formaldehyde were considered (day 0 = autopsy and immersion, five brains, 175 datasets over 0–150 days).

##### General fit

MD and FA were modeled jointly across all brains and time points, using the models of Section 2.5.2, to summarize the overall fixation trend. The eigenvalues *λ* _1–3_ were analyzed analogously (Appendix Figure A1).

##### Brain-specific fits

To capture inter-brain heterogeneity, FA and MD were fitted separately for each brain using median CC values per time point.

##### Harmonizing fit parameters across brains

Where fit parameters were compared across brains, the preferred model could differ between specimens. Because the monoexponential model is the special case *d* = 0, the biexponential definitions apply throughout: total amplitude *A* = *b* + *d*, saturation *S* = *a−b−d*, and effective rate *c*_eff_ = (*|b|c*_1_ + *|d|c*_2_)*/*(*|b|* + *|d|*) with time constant *τ^y^* = 1*/c*_eff_. For a monoexponential fit these reduce to *A* = *b*, *S* = *a−b*, and *τ^y^* = 1*/c*_1_.

#### 2.5.5 Post-mortem interval: temporal evolution fits and tissue condition associations (corpus callosum)

**(i) Temporal evolution during fixation:** For each brain, median CC FA was fitted against time in fixative with the saturation models of Section 2.5.2 (preferred model retained). From the FA fit we obtained FA(*t*_0_), amplitude ΔFA, time constant *τ* (days), and saturation FA, with approximate 95 % intervals. Associations of these quantities with PMI were assessed by ordinary least-squares regression.
**(ii) Discrete tissue conditions vs. PMI:** Using the tissue conditions defined in Section 2.5.1, brain-wise mean CC FA and MD were regressed against PMI by ordinary least squares. Slopes, *p*-values, and *R*^2^ are given in Supporting Information Figure S10.

#### 2.5.6 Temporal evolution fixation effects across anatomical ROIs

For selected anatomical ROIs (WM: corpus callosum and internal capsule, dGM: pallidum, thalamus, putamen, and caudate, cGM: temporal lobe), median FA and MD were computed per acquisition during fixation and plotted over time for three exemplar specimens with dense temporal sampling (brain3, brain6, brain5). This analysis was descriptive and did not include saturation modeling.

#### 2.5.7 Fiber orientation dispersion (***κ***) and fixation-related changes

##### *κ* estimation and stratification

Fiber orientation dispersion (*κ*) was estimated from b-value 4000 s/mm^2^data with the NODDI–DTI implementation in ACID [35] and used to stratify WM voxels. For each brain, a mean *κ* map was computed from the last ten available time points in formaldehyde. WM voxels above a 95 % probability threshold were sorted by *κ* and split into five equally sized groups (G1–G5). For each brain and *κ*-group, mean FA per fixation time point was fitted with the saturation models; model selection followed the AICc/ER procedure described in the Supporting Information (Section 7.5.1). The fitted parameters were amplitude, *τ* (days), and saturation. Model-specific coefficients were harmonized as in Section 2.5.4.

##### Association with fixation kinetics

Fixation-related fit parameters were related to mean group *κ* across all brain–group points using Pearson correlation and ordinary least-squares regression (*Y ∼ κ*).

## 3 Results

### 3.1 Diffusion parameters across tissue conditions

Overall, MD decreased substantially and systematically from in-vivo to in-situ and through subsequent ex-vivo conditions, whereas FA did not show a consistent trend (Figure 2).

For MD, the dominant change occurred between in-vivo and in-situ (relative change of *−*81 %, from 0.728 to 0.140 *×* 10^-3^ mm^2^/s), corresponding to a 5.2-fold reduction before any chemical fixation. The further decrease from beginning of fixation to fixed was comparatively small (relative change of *−*12.5 %, from 0.093 to 0.082 *×* 10^-3^ mm^2^/s), and the total change from in-vivo to the fixed state reached *−*88.8 % (Figure 2 **A**).

FA showed a qualitatively different pattern. Median FA increased from in-vivo to in-situ (relative change of +7.6 %, from 0.605 to 0.651), then decreased after excision and fixation (annotated change at beginning of fixation: *−*11.7 %, Figure 2**B**). The additional change between beginning of fixation and fixed was small (relative change of *−*3.2 %, from 0.535 to 0.518), and the overall change from in-vivo to fixed was *−*14.5 %. The smallest relative parameter shifts in this comparison occurred between beginning of fixation and fixed for both FA and MD.

### 3.2 Protocol comparison using fixation-model fit quality

The diffusion protocol substantially affected the error with which saturation models described fixation-associated MD changes: nRMSE decreased with increasing b-value and was lowest at b-value 4000 s/mm^2^(Figure 3**G**). Although monoexponential and biexponential models showed the same b-value-dependent nRMSE trend, the Supporting Information AICc/ER comparison selected the monoexponential model for MD in the CC across b-values after the parsimony rule (Supporting Information Table S3). In contrast, no comparable protocol dependence was observed for FA: nRMSE remained comparatively high across b-values and did not identify a consistently well-fitting common saturation model (Figure 3**H**; Supporting Information Figure S5). Because MD provided the more reproducible fixation marker and its nRMSE was lowest at b-value 4000 s/mm^2^, this weighting was used for all subsequent main analyses.

Qualitative fixation patterns were shared across weightings: MD decreased toward a plateau, whereas FA remained heterogeneous between brains. Lower b-values yielded higher absolute MD/FA and greater scatter without changing time-course shape. Across tissue conditions, including in-situ, b-value 4000 s/mm^2^showed the lowest voxelwise parameter variability in the CC (Supporting Information Figures S2 and S3).

### 3.3 Temporal evolution fixation dynamics in the corpus callosum

#### Fixation-related changes and model fit for MD

MD decreased rapidly within the first few weeks of fixation and approached a plateau thereafter (Figure 3**E**), with 99.9 % of total MD change within the first 30 days. At b-value 4000 s/mm^2^, the preferred monoexponential model captured this saturation trajectory; alternative biexponential fits were nearly indistinguishable (Supporting Information Table S3). The fitted time constant was *τ ≈* 4–5 days (Section 2.5.2), the time scale on which MD decreased toward its asymptotic fixation value. The amplitude of the decrease was approximately 0.07–0.10 *×* 10*^−^*^3^ mm^2^/s, a reduction of roughly 50–55 % relative to the initial value.

#### Fixation-related changes and model fit for FA

FA showed a less uniform fixation response than MD (Figure 3**F**). Time courses generally decreased or stabilized after an initial transient but did not converge onto a single common pattern across brains, and no single saturation model provided a robust common fit across specimens. This heterogeneity was more pronounced in whole-brain WM, where model selection additionally favored null or simpler models for several protocols (Supporting Information Table S3), motivating the subsequent analyses of post-mortem interval (PMI) (Section 3.4) and within-WM fiber orientation dispersion (*κ*, Section 3.6).

### 3.4 Post-mortem interval (PMI)

#### 3.4.1 PMI dependence of temporal evolution fixation kinetics

As FA did not follow a common saturation trajectory across brains (Section 3.3), we next tested whether brain-specific associations suggested that FA fixation kinetics varied with PMI. In the FA fits (Figure 4**A**), FA decreased over the first *∼*30–50 days toward a plateau. For brain5 (24 h PMI) fixation was best described by a biexponential model (dashed lines), whereas brain3 (15 h PMI), brain6 (18 h PMI) and brain1 (21 h PMI) were best described by a monoexponential fit (solid lines). The initial value FA(*t*_0_) decreased with increasing PMI (Figure 4**B**). The fitted time constant *τ* increased with PMI (Figure 4**D**), indicating slower FA fixation dynamics after longer PMI. By contrast, fitted amplitude ΔFA showed no clear relationship with PMI (Figure 4**C**), and saturation FA only a weak trend (Figure 4**E**).

**Figure 4:**
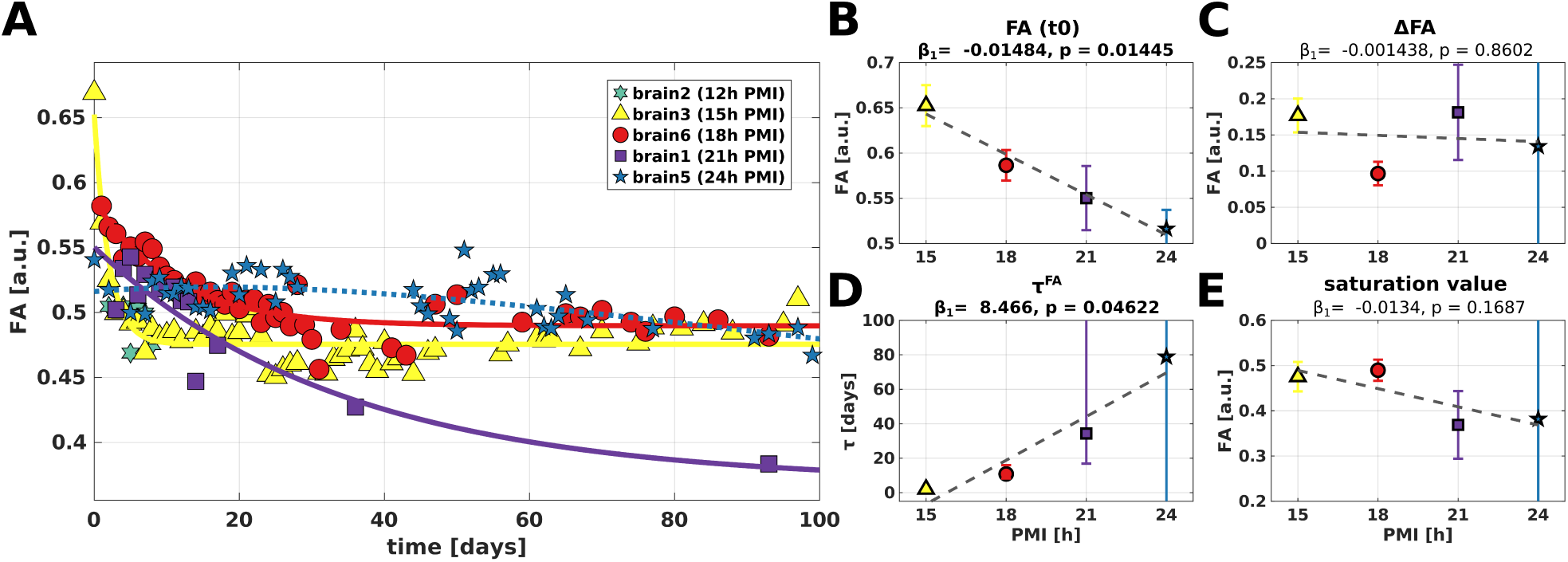
Post-mortem interval (PMI) in relation to temporal evolution during fixation in the corpus callosum (CC): (**A**) Median fractional anisotropy (FA) during immersion fixation (days) for individual brains (distinct markers), with the preferred monoexponential (solid) or biexponential (dashed) fit selected by AICc, evidence ratios, and parsimony (Supporting Information Tables S5 and S6). Ordinary Least Squares (OLS) associations between post-mortem interval (PMI) and fitted FA parameters are shown for (**B**) initial value FA(*t*_0_), (**C**) amplitude ΔFA, (**D**) time constant *τ*, and (**E**) saturation FA. Markers denote individual brains. Vertical error bars indicate approximate 95 % confidence intervals of the fitted parameters. Dashed gray lines show the OLS regression across brains. Inset statistics report the OLS slope (*β*_1_) and *p*-value. Saturation modeling and the PMI–parameter associations in (**B**)–(**E**) included four brains (brain3, brain6, brain1, and brain5). Brain2 contributed too few fixation time points for reliable model fitting and is therefore absent from (**B**)–(**E**). Complete OLS and Pearson statistics are provided in Supporting Information Table S7.

### 3.5 Temporal evolution fixation effects across anatomical ROIs

FA showed a consistent dissociation between WM and grey matter (GM) across the three specimens (PMI: 15, 18, and 24 h). In the corpus callosum and internal capsule, FA decreased during days 0– 10 (median relative decrease *∼*15 %), whereas gray-matter ROIs showed modest early increases or upward drifts (median relative change up to *∼*8 %). Thus, WM and GM changed in opposite directions before stabilizing (Figure 5**B**–**D**). Within GM, this pattern was similar in regions differing in depth and proximity to external or ventricular surfaces: the temporal lobe near the external surface, the thalamus bordering ventricular cerebrospinal fluid (CSF), and the putamen deeper without direct CSF contact. Their similar time courses showed no consistent ordering by anatomical depth. Differences among dGM regions were modest, although pallidum and thalamus changed slightly more than putamen and caudate nucleus. Regional FA differences were less pronounced in brain5 (24 h PMI) than in brain3 and brain6 (15–18 h PMI).

**Figure 5:**
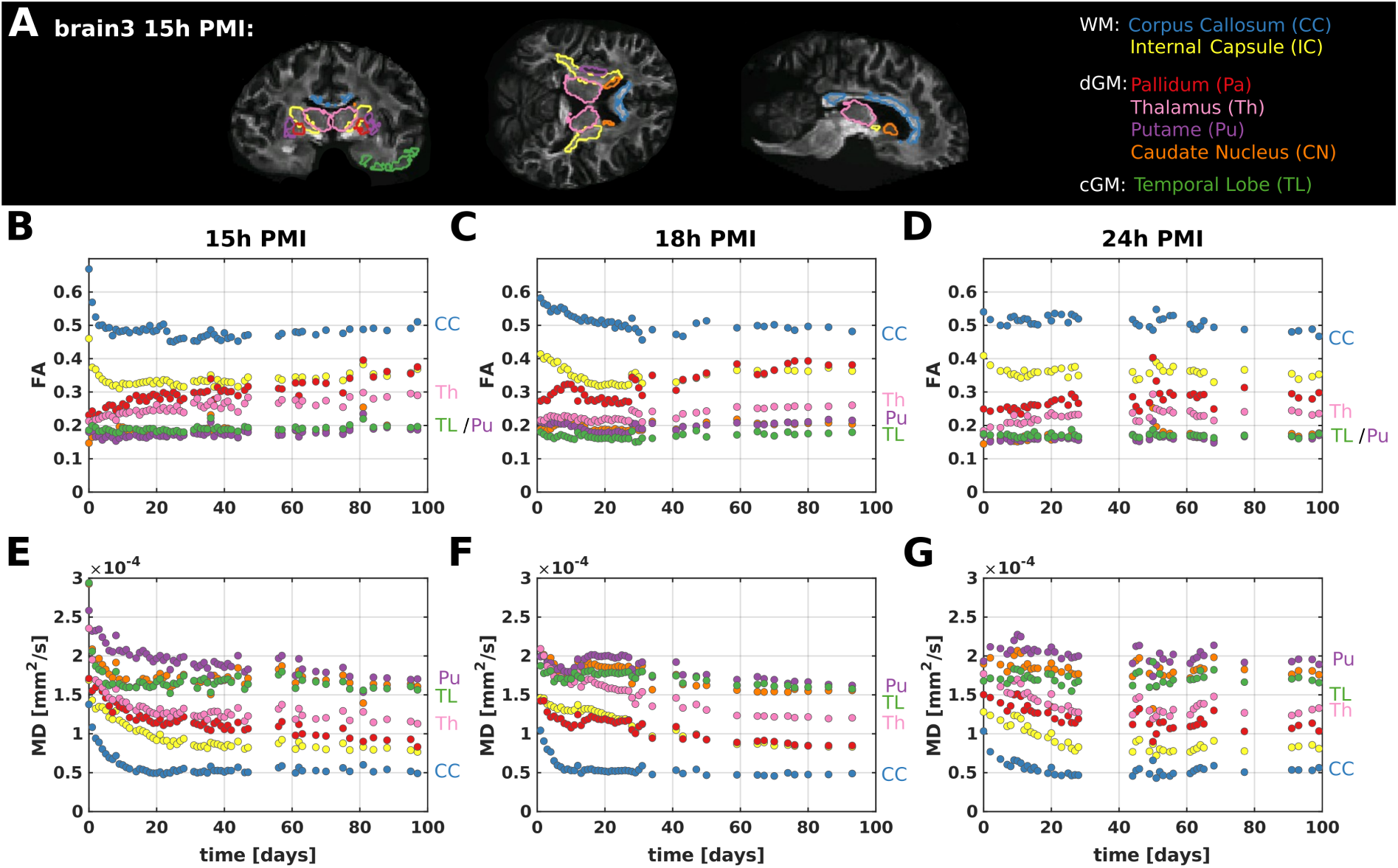
Temporal evolution of FA and MD during fixation across anatomical ROIs: (**A**) Exemplary region of interest outlines for brain3 (15 h PMI) on coronal, axial, and sagittal slices, with region of interest (ROI)s: white matter (WM) (corpus callosum, internal capsule), deep grey matter (dGM) (pallidum, thalamus, putamen, caudate nucleus), and cortical grey matter (cGM) (temporal lobe). (**B**)–(**D**) Median fractional anisotropy (FA) over fixation time (days) for brain3 (15 h PMI), brain6 (18 h PMI), and brain5 (24 h PMI, columns from left to right). (**E**)–(**G**) Median mean diffusivity (MD) over the same fixation period for the same specimens. Colors encode ROIs as in (**A**).

MD showed similar kinetics for all regions: an initial decrease over the first 10 days (mean relative change *−*19 %), followed by a plateau after approximately 20 days (Figure 5**E**–**G**). Absolute MD remained stratified by tissue class throughout fixation, lowest in WM and higher in gray matter.

### 3.6 Fiber orientation dispersion (***κ***) and fixation-related changes

To test whether fixation-related FA changes within WM relate to local fiber organization, voxels were stratified into five *κ* groups (Figure 6**A**). Across all three brains, fitted FA amplitude ΔFA depended on *κ*: lower-*κ* groups showed weaker fixation-related changes or modest early decreases, whereas higher-*κ* groups showed stronger upward drifts during fixation (Figure 6**B**–**D**).

**Figure 6:**
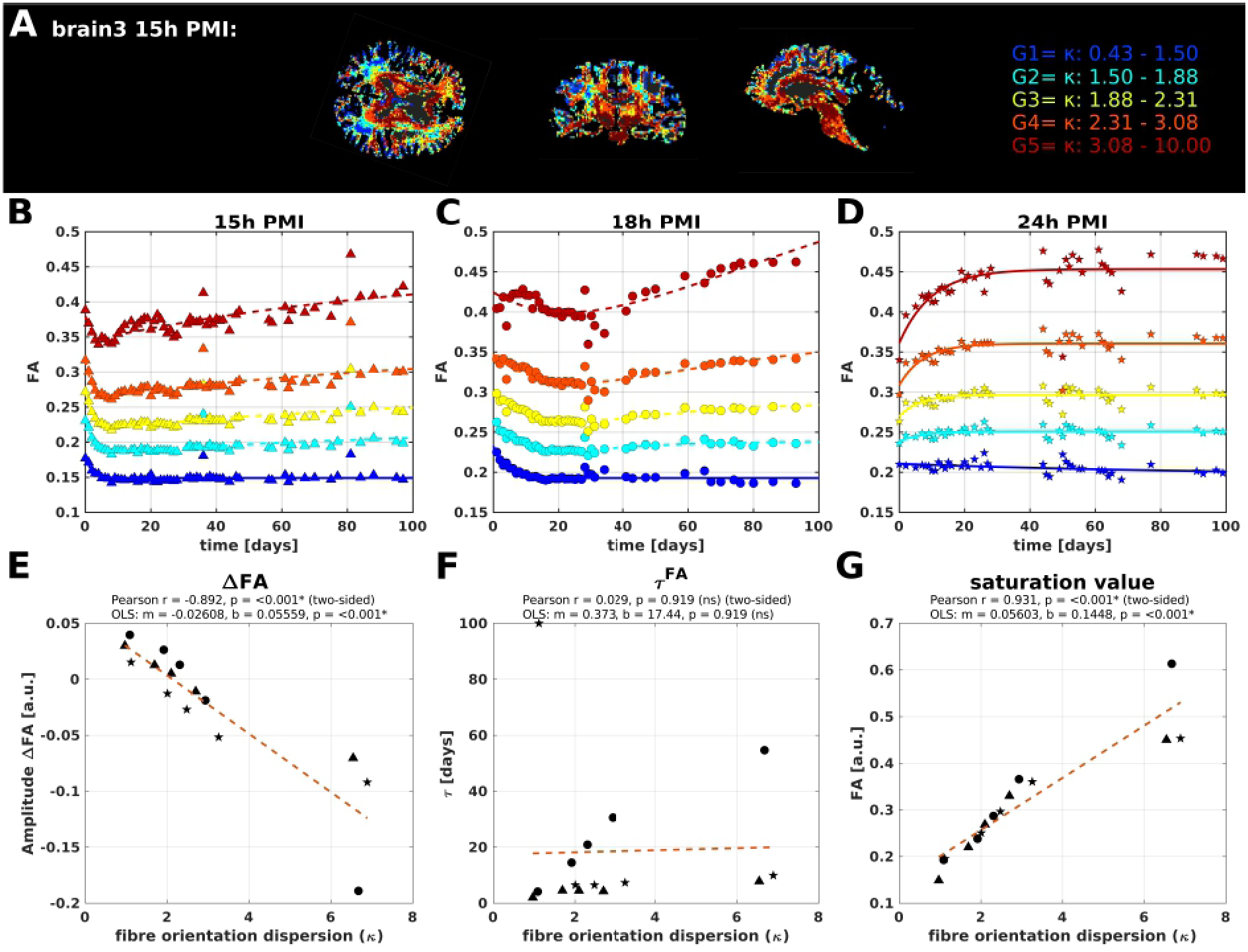
Influence of fiber orientation dispersion (*κ*) on temporal evolution of fractional anisotropy (FA) during fixation in white matter (WM): (**A**) Representative slices (brain3, 15 h PMI) with voxels assigned to five disjoint *κ* groups G1–G5. (**B**)–(**D**) Mean FA over fixation time for G1–G5 for PMI 15 h, 18 h, and 24 h, respectively. Solid lines show monoexponential fits and dashed lines show biexponential fits, selected per group by ER. (**E**)–(**G**) Associations between mean group *κ* and fitted FA amplitude ΔFA, *τ*, and saturation FA. Each marker denotes one brain–group combination. Statistics in each panel report Pearson correlation and OLS regression. The dashed orange line shows the OLS fit. Complementary mixed-effects analyses accounting for brain as a random effect are provided in the Supporting Information (Section 7.8).

Quantitatively, the fitted amplitude became more negative with increasing mean *κ* (Pearson *r* = *−*0.892, *p <* 0.001; OLS slope *m* = *−*0.0261, *p <* 0.001), indicating a stronger fixation-related FA change in more coherently organized fiber regions. The time constant *τ* showed no consistent association with mean *κ* (Pearson *r* = 0.029, *p* = 0.919; Figure 6**F**). Saturation FA also increased with mean *κ* (Pearson *r* = 0.931, *p <* 0.001; OLS slope *m* = 0.0560, *p <* 0.001). Fiber orientation dispersion was therefore associated with the magnitude and asymptotic level of fixation-related FA changes, rather than their temporal scale, consistent with the eigenvalue-based analysis in the Appendix (Figure A1).

## 4 Discussion

Post-mortem changes in dMRI of the human brain across tissue conditions and during fixation affect mean diffusivity (MD) and fractional anisotropy (FA) differently. Interpreting these changes therefore requires separating early post-mortem cellular and biophysical changes from later chemical fixation and accounting for diffusion weighting and local microstructure.

### 4.1 Tissue condition transitions and fixation-time dependence reveal distinct effects on MD and FA

Section 3.1 shows that MD and FA are affected by different parts of the post-mortem process. The dominant MD shift occurred between in-vivo and in-situ, before brain extraction and chemical fixation, indicating that the largest diffusivity reduction is driven mainly by acute changes after circulatory arrest rather than fixation chemistry.

Although this MD shift preceded chemical fixation, immersion fixation still produced a largely mono-exponential MD decrease toward a plateau (Section 3.3); dense early sampling made this later fixation effect detectable here.

FA followed a different pattern (Section 3.1), with a comparatively small in-vivo-to-in-situ change but larger shifts after excision and during fixation. The in-vivo-to-in-situ comparison should be interpreted in the context of two unavoidable confounds: cohort age and tissue temperature differed between the in-vivo reference and the post-mortem donor scans. Sensitivity bounds in the Supporting Information (Section 7.2; Supporting Information Table S1) indicate that neither age nor cooling can explain the large in-vivo-to-in-situ MD decrease; using Berger post-mortem temperature slopes, cooling may account for only a small fraction of its magnitude (*∼*11 %), whereas age predicts the opposite direction. Protocol mismatch is a further caveat for that comparison (in-vivo *b* = 1000 vs. ex-vivo *b* = 4000 s/mm^2^): within the same post-mortem tissue, switching from *b* = 1000 to *b* = 4000 lowers median FA by *∼*7.5–14.5 % and absolute MD (in-situ medians *∼*0.22 vs. *∼*0.15 *×* 10^-3^ mm^2^/s; Supporting Information Figure S3), an offset far smaller than the observed MD drop (*−*81 %) and opposite in sign to the observed FA increase (+7.6 %). For FA, age and temperature can affect absolute values, but the estimated effects do not fully explain the observed difference; the within-cohort in-situ-to-beginning-of-fixation comparison therefore provides the stronger evidence that FA is sensitive to later tissue handling and fixation. Strongest post-mortem inferences overall rest on repeated measurements within the same ex-vivo brains.

### 4.2 Temporal modeling during fixation: MD as a robust fixation marker and FA as a microstructure-sensitive diffusion parameter

During fixation, MD and *λ*_1–3_ decreased in whole-brain WM and all four parameters were well described by an exponential saturation (Appendix Figure A1). The MD saturation trajectory was reproducible across brains, whereas FA was more heterogeneous and requires microstructure-aware interpretation (Section 3.3).

The time course is consistent with progressive formaldehyde cross-linking [25], with most MD change within the first 30 days of immersion (Section 3.3), in line with fixed porcine tissue [29] and *R^∗^* in the same dataset [38]. A relaxation-mediated contribution is plausible via coupling between compartmental relaxation and MD [41, 42].

The tensor response during fixation was not isotropic. The Appendix analysis showed a stronger decrease in *λ*_1_ than in *λ*_2_ and *λ*_3_, indicating stronger effects along the principal fiber direction than perpendicular diffusion and providing a deeper basis for understanding the observed heterogeneity of the FA changes. Even for MD, the fixation response is shaped by tissue architecture rather than a purely scalar reduction in water mobility.

In contrast to MD, FA did not follow a single common fixation-time trajectory across brains (Section 3.3). This heterogeneity covaried with PMI, regional anatomy, and fiber organization (Section 4.3) and was strongest in highly organized WM.

### 4.3 Sources of heterogeneity in fixation behavior: PMI, regional anatomy, and fiber organization

Fixation heterogeneity, particularly for FA, is not explained by time in fixative alone, but is associated with PMI, regional anatomy, and fiber organization.

PMI can alter tissue state and subsequent fixative response through autolysis-related changes such as myelin decomposition [43, 44, 45, 30]. Here, PMI was related not only to discrete tissue conditions but also to brain-specific fixation-time parameters (Section 3.4).

The fitted parameters suggest an association between PMI and tissue state at fixation onset and subsequent trajectory pace. At shorter PMIs (15–18 h), regional fixation trends remained more distinct (Figure 5), whereas at 24 h regional FA differences were less pronounced. Absolute PMI effects within 12–24 h remained small relative to principal tissue condition shifts; at discrete conditions, MD showed a clear association with PMI at beginning of fixation and in the fixed state, and FA after hydration, whereas PMI-related FA effects were also apparent in fitted fixation parameters, particularly FA(*t*_0_) and *τ* (Section 3.4).

Regional and *κ*-based analyses (Sections 3.5 and 3.6) indicate that fixation behavior cannot be explained by anatomical depth alone. The regional data showed no simple ordering by proximity to an external or ventricular surface; the clearest distinction was between WM and gray matter (Section 3.5).

Across scales, the ROI, *κ*, and eigenvalue analyses converged on the same pattern: the more strongly directed the tissue organization, the stronger the fixation-related effect, whether comparing WM with gray matter, higher-with lower-*κ* WM, or *λ*_1_ with *λ*_2_ and *λ*_3_.

Main *κ* associations used Pearson correlation and OLS regression and thus did not model brain as a grouping variable; complementary linear mixed-effects model (LME) analyses with brain as a random intercept confirmed the same principal FA pattern and also showed *κ* associations for all three MD fixation parameters (Supporting Information, Section 7.8). Because *κ* is estimated from FA in the NODDI– DTI framework [41], the relation between the saturation value of FA and *κ* is expected (Figure 6), whereas the correlations of ΔFA, ΔMD, and *τ*^MD^ with *κ* are not predicted by that dependence alone (Supporting Information, Section 7.8). Although *κ* is not an independent histological ground truth, it remains the most direct MRI-based proxy for local fiber alignment here and a useful explanatory variable for testing whether fixation behavior follows tissue organization rather than geometric depth alone.

### 4.4 Protocol optimization and implications for post-mortem diffusion MRI

Protocol comparison (Section 3.2) shows that fixation-related changes depend on diffusion weighting as well as tissue biology. Here, b-value 4000 s/mm^2^ minimized parameter variance across brains and tissue conditions, extending porcine work [29] to whole human brains from in-situ through late immersion. If the recommendation that the product of MD and b-value should be *∼*1 [19] were taken literally, this would imply b-values larger than 10000 s/mm^2^ for post-mortem human brain tissue. Our fixed ex-vivo MD in WM was smaller than 0.1 *×* 10*^−^*^3^ mm^2^/s, corresponding to a factor of *∼*10 relative to in-vivo-like weighting rather than the factor of *∼*4 reported by Dyrby et al. [29] for porcine tissue with diffusivity *∼*0.2 *×* 10^-3^ mm^2^/s. Contributing reasons for the smaller MD include (i) the higher b-value itself (Supporting Information Figure S3), (ii) temperature [32], and (iii) PMI (Figure S10).

The reduced variance of the fixation trajectories at higher weighting is particularly informative: it suggests that higher diffusion weighting better captures subtle fixation-related microstructural effects, rather than merely yielding numerically smoother estimates, and provides the strongest indication in this study that higher b-values can improve precision for detecting subtle microstructural differences in post-mortem tissue, potentially beyond fixation alone. The finding that b-values below 4000 s/mm^2^ lead to increased variance may also help explain why fixation-related changes can be difficult to detect in studies using substantially lower b-values, such as Shatil et al. [28] (b-value 700 s/mm^2^). Although signal-fit root mean square error (RMSE) is expected to depend on b-value, the two matched b-value 2000 s/mm^2^ datasets still differed in RMSE, indicating that residual protocol factors also affected signal-fit quality (Supporting Information, Section 7.4).

### 4.5 Limitations

We analyzed six whole-brain ex-vivo specimens, five of which contributed longitudinal formaldehyde fixation data. Although, to our knowledge, this is one of the most densely sampled longitudinal whole-brain ex-vivo dMRI datasets to date, the sample remains small and results warrant caution. Nevertheless, the cohort sufficed to resolve reproducible MD saturation during fixation, compare diffusion weighting across tissue conditions, and characterize main FA heterogeneity sources.

Diffusion weightings (1000–4000 s/mm^2^) differed beyond b-value in TE/TR, distortion correction, coil, and, for one brain, resolution (Table 1), so this was not a pure b-value experiment. Matched 2000 s/mm^2^ shells nevertheless supported diffusion weighting as the dominant factor for absolute MD/FA offsets (Section 7.4). We did not aim to define a universal b-value; b-value 4000 s/mm^2^ operationally reduced fit error and variance here, whereas similar fixation trends persisted at b-value 1000 s/mm^2^ with lower precision (Section 3.2).

Scanning temperature was not measured, though brains were scanner-room temperature acclimated. Temperature may affect absolute MD and FA [34, 32, 33], but Supporting Information sensitivity bounds indicate that residual room-temperature variation cannot explain the fixation-related changes alone (Supporting Information, Section 7.2; Supporting Information Table S1). PMI and uneven tissue-condition coverage remain incompletely characterized for whole human brains and may contribute particularly to FA heterogeneity, though within 12–24 h their impact remained small relative to principal tissue-condition shifts. Without histological validation of fixative penetration or microstructural change, associations with tissue class, regional anatomy, *κ*, and tensor eigenvalues should be read as MRI-based rather than histologically proven.

Longitudinal coregistration and tissue segmentation were prerequisites for regional and *κ* analyses; imprecision may add trajectory noise. Future work should test generalisability across diffusion models, platforms, partial brain samples, fixatives, and temperature-controlled or histology-linked designs modeling PMI and fixative penetration.

## 5 Conclusion

Post-mortem processes in whole human brains such as early post-mortem physiology and subsequent chemical fixation contribute differently to MD and FA. Dense longitudinal sampling across in-situ, fixation, and hydration conditions showed that the largest MD reduction occurred between the in-vivo reference and in-situ tissue, before fixation, followed by a reproducible monoexponential MD saturation during immersion fixation. FA changed more after brain extraction and early fixation, but without a common fixation related trajectory across specimens. That heterogeneity covaried with PMI, regional anatomy, and within-WM fiber orientation dispersion (*κ*).

Among the tested protocols, b-value 4000 s/mm^2^ minimized parameter variance across tissue conditions. Lower weightings preserved the same qualitative MD and FA trajectories, with higher absolute estimates and reduced precision. Under the present acquisition setting, MD is therefore a comparatively reproducible temporal marker of immersion fixation, whereas FA should not be used as a fixation clock and requires a microstructure-aware interpretation across tissue conditions, regions, and specimens. The higher weighting is operationally preferable for precision here, and is not claimed as a universally optimal ex-vivo b-value.

## Acknowledgments

This work was supported by ERA-NET NEURON (hMRI-ofSCI); the German Federal Ministry of Education and Research (BMBF; 01EW1711A and 01EW1711B); the Forschungszentrum Medizintechnik Hamburg (fmthh; grant 01fmthh2017); and the Deutsche Forschungsgemeinschaft (DFG, German Research Foundation), including the Emmy Noether Programme (MO 2397/4-1 and MO 2397/4-2), DFG Priority Program 2041 “Computational Connectomics” (MO 2397/5-1, MO 2397/5-2, MO 2249/3-1, and MO 2249/3-2), and project number 347592254 (WE 5046/4-2). Additional support was provided by the European Research Council under the European Union’s Seventh Framework Programme (FP7/2007–2013) / ERC grant agreement no. 616905, and by ERC grant MRStain (grant agreement ID: 101089218; DOI: 10.3030/101089218). Funded by the European Union. Views and opinions expressed are those of the author(s) only and do not necessarily reflect those of the European Union or the European Research Council Executive Agency. Neither the European Union nor the granting authority can be held responsible for them.

AI-assisted tools were used solely for spelling correction and for shortening or tightening the wording of the manuscript. They were not used to generate scientific content, figures, or analyses.

## Data Availability Statement

Analysis code supporting this study is being prepared and will be made available in a public GitHub repository. The imaging data that support the findings of this study are available from the corresponding author upon request.

## 6. Appendix

### 6.1 Methods

**Table A1:**
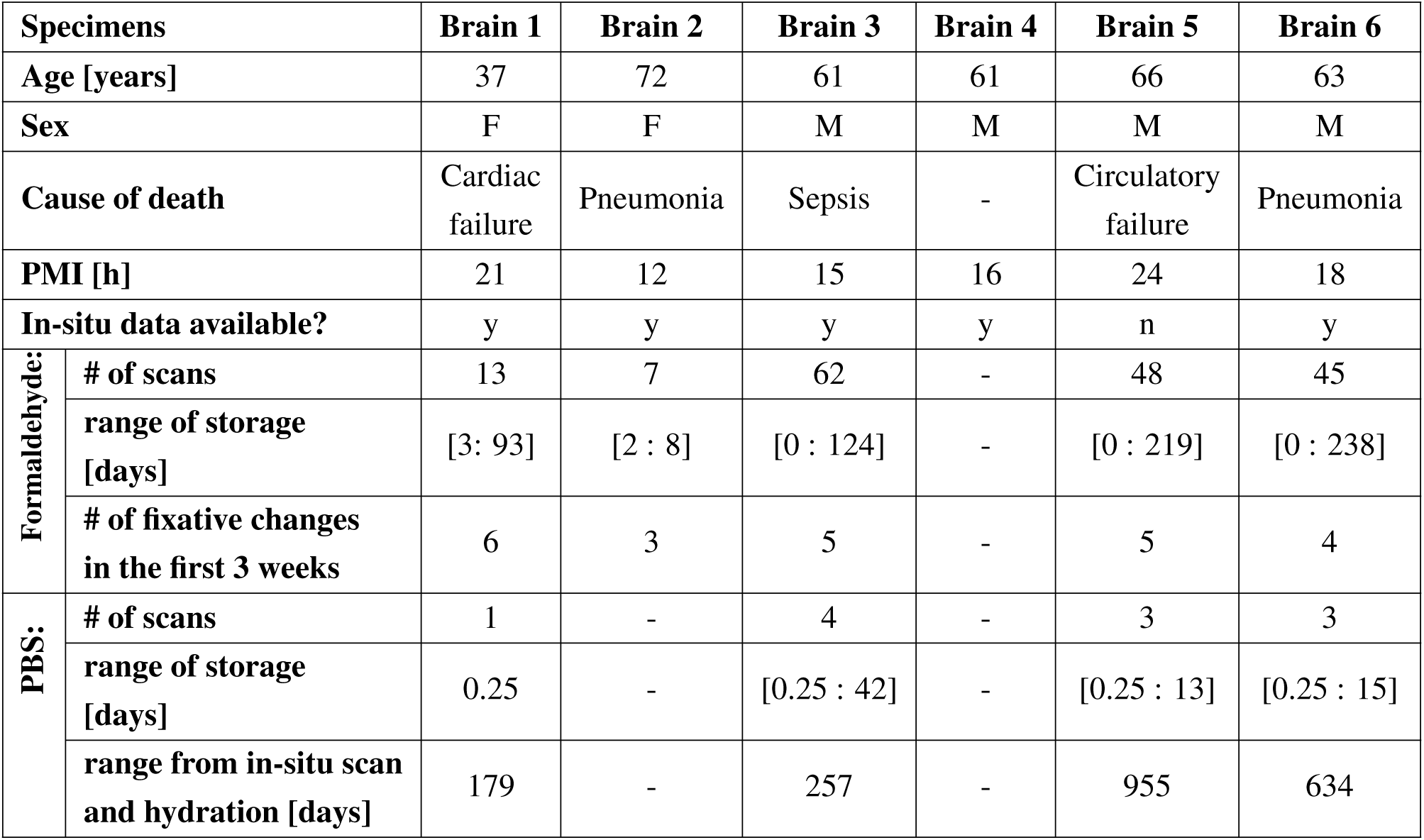
Detailed information of the whole human brains (brain 1–6), including scan availability and storage information for the different tissue conditions. Fixation time points refer to MRI measurements acquired within the first 150 days after immersion in formaldehyde. For the formaldehyde and PBS rows, # of scans gives the number of available MRI measurements in the respective medium, and range of storage denotes the span of days on which MRI measurements were acquired in that medium.

| Specimens |  | Brain 1 | Brain 2 | Brain 3 | Brain 4 | Brain 5 | Brain 6 |
| --- | --- | --- | --- | --- | --- | --- | --- |
| Age [years] |  | 37 | 72 | 61 | 61 | 66 | 63 |
| Sex |  | F | F | M | M | M | M |
| Cause of death |  | Cardiac failure | Pneumonia | Sepsis | - | Circulatory failure | Pneumonia |
| PMI [h] |  | 21 | 12 | 15 | 16 | 24 | 18 |
| In-situ data available? |  | y | y | y | y | n | y |
| Formaldehyde: | # of scans | 13 | 7 | 62 | - | 48 | 45 |
|  | range of storage [days] | [3: 93] | [2 : 8] | [0 : 124] | - | [0 : 219] | [0 : 238] |
|  | # of fixative changes in the first 3 weeks | 6 | 3 | 5 | - | 5 | 4 |
| PBS: | # of scans | 1 | - | 4 | - | 3 | 3 |
|  | range of storage [days] | 0.25 | - | [0.25 : 42] | - | [0.25 : 13] | [0.25 : 15] |
|  | range from in-situ scan and hydration [days] | 179 | - | 257 | - | 955 | 634 |

### 6.2 Modeling the Exponential Decay of Diffusion Parameters during Fixation

**Figure A1:**
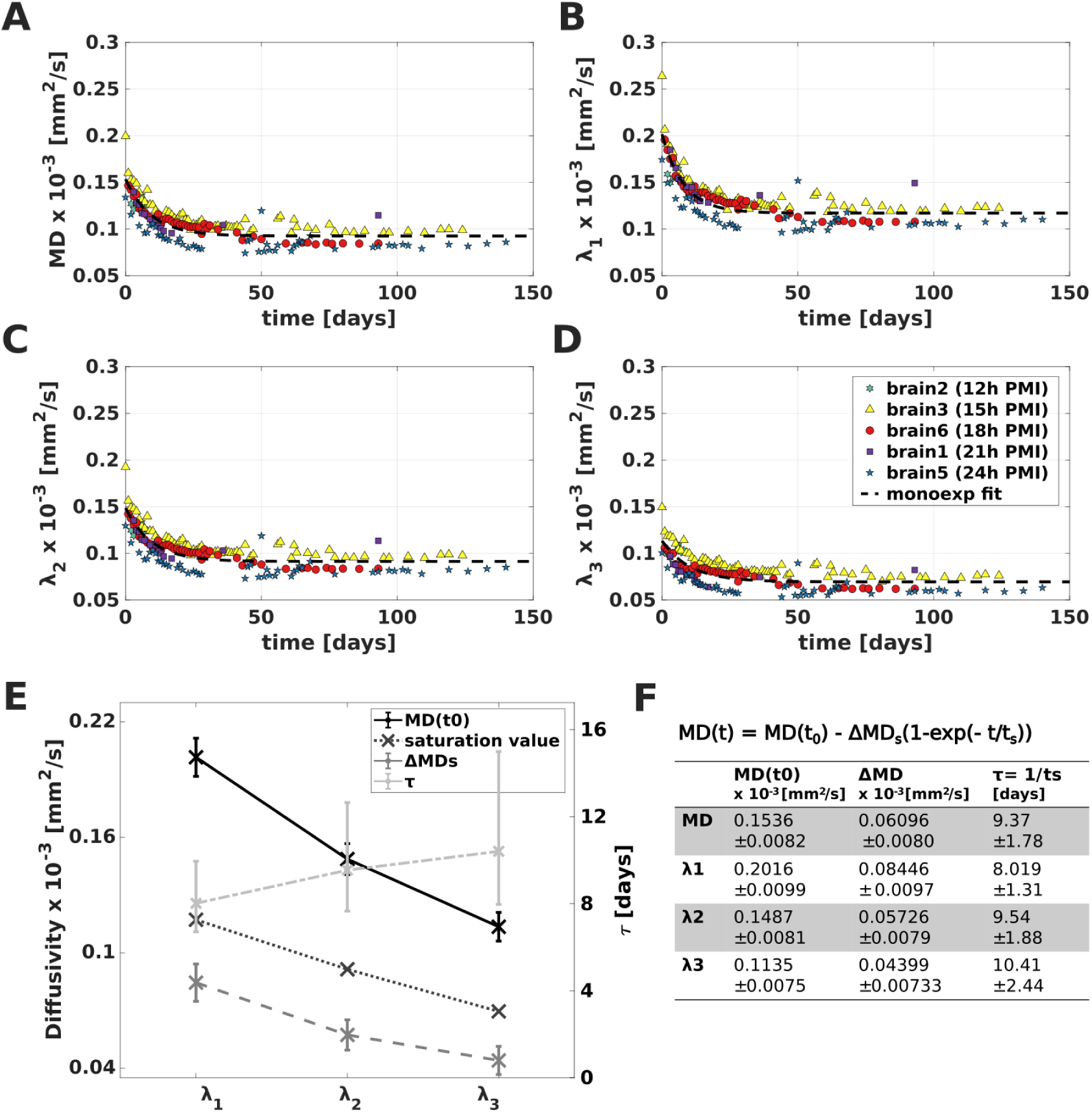
Temporal evolution of diffusion parameters during fixation in the WM region: Plots of MD (**A**) and the eigenvalues *λ*_1-3_ (**B-D**) estimated using the DTI model of the diffusion-weighted (dw) images at b=4000 s/mm^2^ over fixation time for different brain specimens in the WM. The colors represent different brain specimens, while the black dashed line represents the selected monoexponential model fit over all data points from all brains. Fitted parameters (with 95 % confidence bounds) for MD and *λ*_1-3_ are plotted against the three tensor eigenvalues *λ*_1-3_, as well as the saturation value (diffusivity after 150 days in fixative) in **E)** and shown in the table (**F)**. The error bars in **E)** represent the 95 % confidence interval. MD(t0) is the initial MD, *Δ*MD is the change in MD over the fixation time, and *τ* is the saturation time constant. The same definitions for MD(t0), *Δ*MD, and *τ* also apply to the three eigenvalues *λ*_1-3_.

All four diffusion parameters decreased during fixation (0–150 days) and were best described by monoexponential fits (Figure A1). *λ*_1_ showed the largest decrease, followed by *λ*_2_ and *λ*_3_; after 30 days, diffusivity had decreased by *∼*35–38 % relative to the first measurement and was within *∼*3 % of the saturation value.

### Abbreviations and acronyms

CC: Corpus callosum
cGM: Cortical grey matter
CSF: Cerebrospinal fluid
dGM: Deep grey matter
dMRI: Diffusion Magnetic Resonance Imaging
DTI: Diffusion tensor imaging
dw: Diffusion-weighted
ECMOCO: Eddy current and motion correction
FA: Fractional anisotropy
FOV: Field of view
GM: Grey matter
HySCO: Hyperelastic susceptibility artifact correction
IQR: Interquartile range
LME: Linear mixed-effects model
MD: Mean diffusivity
MNI: MNI space
MPM: Multi-parametric mapping
MR: Magnetic resonance
MRI: Magnetic resonance imaging
NLLS: Non-linear Least Squares
nRMSE: Normalised root mean square error
OLS: Ordinary Least Squares
PBS: Phosphate-buffered saline
PMI: Post-mortem interval
RMSE: Root mean square error
ROI: Region of interest
SNR: Signal-to-noise ratio
TE: Echo time
TPM: Tissue probability map
TR: Repetition time
WM: White matter

## Supporting Information contents

*The following sections are provided as Supporting Information for review and online publication*.

*—* Fixation and hydration solutions
*—* Sensitivity bounds for temperature and age effects
*—* Registration and Segmentation Pipeline

**–** Used time intervals for analysis using tissue conditions
*—* Protocol Comparison

**–** Methods: Comparative study of diffusion protocols
**–** Supplementary Results: Comparative study of diffusion protocols
**–** Protocol comparison in corpus callosum and whole white matter
*—* Modeling the temporal evolution of diffusion parameters during fixation and protocol comparison

**–** Model selection and evidence ratios
*—* Supplementary statistics: PMI associations with fixation-model parameters
*—* Supplementary statistics: PMI associations at discrete tissue conditions

**–** PMI dependence across discrete tissue conditions

*—* Supplementary analysis: fiber orientation dispersion and fixation parameters

