## Supplementary Material for "Diffusion tensor imaging of whole human brains during long-term formaldehyde fixation: Temporal evolution of diffusion parameters, post-mortem conditions, and dependence on tissue structure"

### 7 Supporting Information

#### 7.1 Fixation and hydration solutions

Immersion fixation and subsequent rehydration used phosphate-buffered saline (PBS) and a 4 % formaldehyde solution in PBS. Solutions were prepared as follows.

**PBS (rehydration and diluent for fixation):** Per litre of aqueous solution:

- disodium hydrogen phosphate ($Na_2HPO_4$): 1.43 g (Roth 4984)
- sodium dihydrogen phosphate ($NaH_2PO_4$): 0.43 g (Roth 2370)
- sodium chloride (NaCl): 7.20 g (Roth 3957)

The pH was adjusted to 7.4.

**4 % formaldehyde fixation solution:** Acid-free 37 % formaldehyde solution for histology (Roth P733.2) was diluted with PBS to a final formaldehyde concentration of 4 %. The pH was checked after dilution.

#### 7.2 Sensitivity bounds for temperature and age effects

Diffusion parameters are sensitive to physical and biological factors that differed between measurements, particularly tissue temperature and cohort age. Because tissue temperature was not measured during scanning, and because the in-vivo reference cohort was younger than the post-mortem donor cohort, we estimated the expected magnitude of these two potential confounds from external reference data. The aim of this analysis was not to correct the data retrospectively, but to test whether temperature or age alone could plausibly account for the observed in-vivo-to-post-mortem and fixation-related effects. The main comparisons are summarized in Table S1.

**Temperature** All specimens were acclimated to scanner-room temperature for several hours before scanning. Because absolute tissue temperature was not measured, we estimated expected temperature effects from Berger et al. [32] and compared them with the effects observed in this study. Berger et al. [32] reported separate WM linear models for MD and FA fitted to post-mortem data only and to data that also included in-vivo measurements. Because the with-in-vivo fit conflates temperature with other living-to-dead confounds, we used the post-mortem-only (ex-vivo) white-matter slopes for both questions below.

**Fixation/ex-vivo effect:** To test whether fixation-related changes could be explained by temperature alone, we considered a conservative residual room-temperature range of 22–26 °C ($\Delta T = 4$ °C) and applied the post-mortem-only white-matter model from Berger et al. [32]. This predicts an expected MD change of $0.017 \times 10^{-3}$ mm$^2$/s (slope-error range: 0.006–$0.029 \times 10^{-3}$ mm$^2$/s) and an expected FA change of 0.0003 a.u. (Berger et al.: not statistically significant). Compared with the modeled fixation-related MD amplitude ($\sim$0.07–$0.10 \times 10^{-3}$ mm$^2$/s) and the early WM FA change of $\pm 0.1$, residual temperature mismatch may add measurement-to-measurement scatter but cannot explain the systematic fixation effects on its own.

**In-vivo to in-situ difference:** To test whether the in-vivo-to-in-situ parameter difference could be explained by temperature alone, we considered cooling from 37 °C to 22 °C ($\Delta T = -15$ °C) and applied the same post-mortem-only (ex-vivo) white-matter slopes from Berger et al. [32]. This predicts an MD decrease of $0.065 \times 10^{-3}$ mm$^2$/s (slope-error range: $0.021$–$0.108 \times 10^{-3}$ mm$^2$/s) and an FA change of $-0.001$ a.u. (not statistically significant). The observed in-vivo-to-in-situ MD decrease ($0.588 \times 10^{-3}$ mm$^2$/s; $-81$ %) is therefore substantially larger than the temperature-only estimate ($\sim$11 % of the observed drop). For FA, we observed an increase from 0.605 to 0.651 (+7.6 %), whereas the Berger model predicts a near-zero decrease on cooling. Temperature alone cannot explain the primary in-vivo-to-in-situ FA difference.

**Age** The in-vivo reference cohort was younger than the post-mortem donor cohort ($35.75 \pm 9$ years versus $60.0 \pm 11$ years). We therefore estimated whether normal age-related white-matter change could account for the observed in-vivo-to-in-situ differences. As an external reference, we used the lifespan white-matter brain-chart data of Kim et al. [46]. Across the adult age range (18–80 years), a quadratic approximation of the Kim et al. data predicted that increasing age from 35.75 to 60 years would decrease FA by approximately 0.017–0.028 a.u. in the corpus callosum and increase MD by approximately 0.037–0.045 $\times$ 10$^{-3}$ mm$^2$/s. A direct median comparison between age windows around the two cohort means gave the same qualitative direction (lower FA and higher MD at older age).

These expected ageing effects cannot explain the primary in-vivo-to-in-situ findings in our data. For MD, ageing predicts a small increase, whereas we observed a large decrease from 0.728 to $0.140 \times 10^{-3}$ mm$^2$/s ($-81$ %). For FA, ageing predicts a decrease, whereas we observed an increase from 0.605 to 0.651 (+7.6 %). Thus, the cohort age difference has the wrong direction for both primary in-vivo-to-in-situ effects and cannot account for the observed post-mortem diffusion changes.

| Potential confound | Expected effect from external reference | Observed effect in this study | Interpretation |
|---|---|---|---|
| Residual temperature during ex-vivo/fixation scans (22–26 °C) | MD: 0.017 $\times$ $10^{-3}$ mm$^2$/s (Berger post-mortem-only model). FA: 0.0003 a.u. | MD fixation amplitude: $\sim$0.07–0.10 $\times$ $10^{-3}$ mm$^2$/s. Early WM FA change: $\pm$ 0.1 a.u. | Residual temperature mismatch may add scatter, but cannot explain the systematic fixation effects on its own. |
| Physiological-to-room-temperature difference for in-vivo-to-in-situ comparison (37–22 °C) | MD: expected decrease of 0.065 $\times$ $10^{-3}$ mm$^2$/s (Berger post-mortem-only/ex-vivo model). FA: $-0.001$ a.u. | MD: $-0.588$ $\times$ $10^{-3}$ mm$^2$/s ($-81$ %). FA: $+0.046$ a.u. ($+7.6$ %). | Temperature may explain only a small fraction of the MD decrease ($\sim$11 %), not its full magnitude. The expected FA effect is negligible and has the wrong direction. |
| Age difference between cohorts (35.75 to 60 years) | MD: expected increase of 0.037–0.045 $\times$ $10^{-3}$ mm$^2$/s from Kim et al. corpus-callosum estimates. FA: expected decrease of 0.017–0.028 a.u. | MD: $-0.588$ $\times$ $10^{-3}$ mm$^2$/s ($-81$ %). FA: $+0.046$ a.u. ($+7.6$ %). | Age has the wrong direction for both primary in-vivo-to-in-situ effects and cannot account for the observed post-mortem changes. |

**Table S1: Summary of temperature and age sensitivity bounds.** Expected effects are external-reference estimates for white matter or corpus-callosum-relevant regions and are compared with the observed in-vivo-to-in-situ or fixation-related effects in this study.

#### 7.3 Registration and Segmentation Pipeline

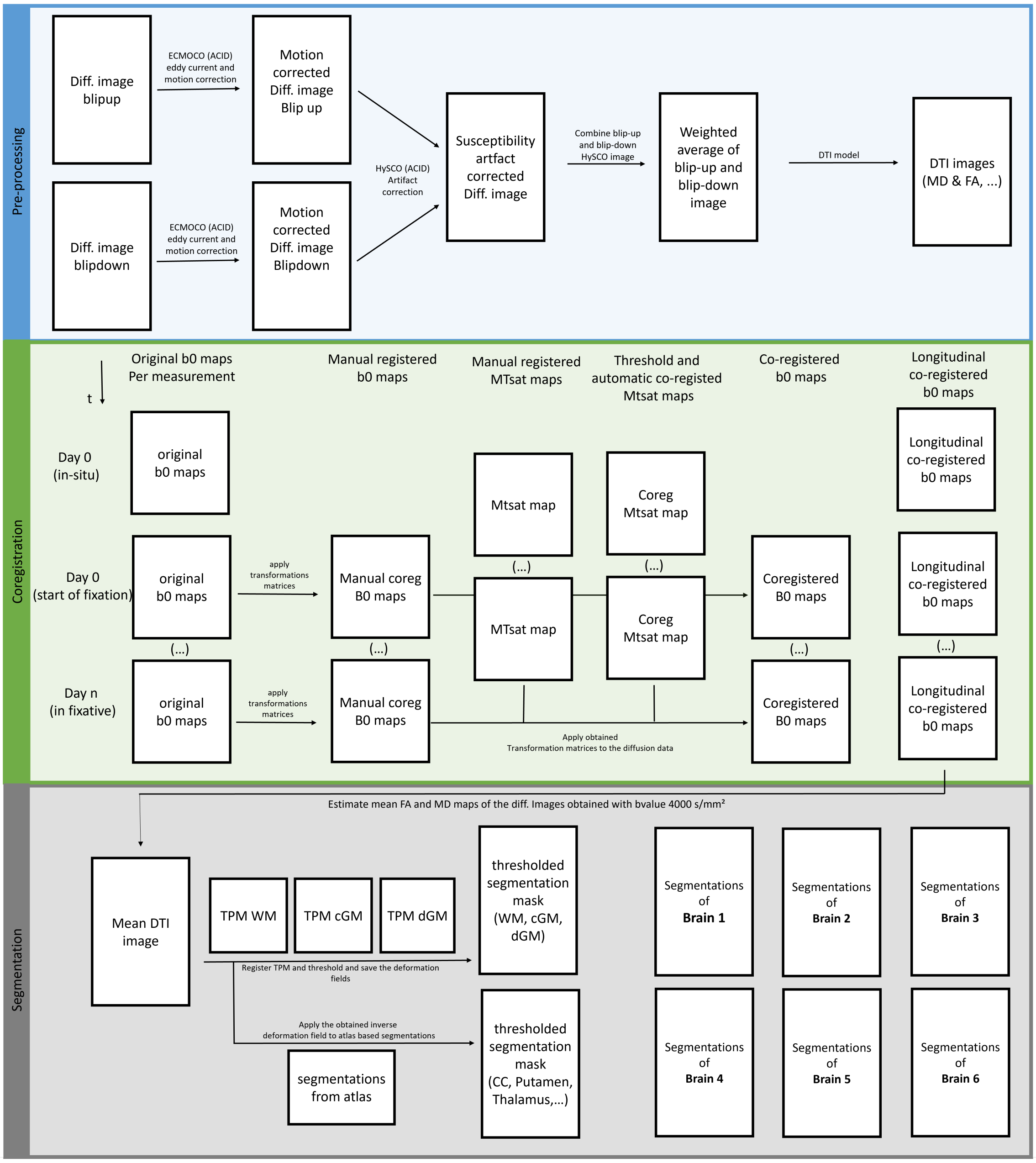


**Figure S1: Overview of the diffusion-MRI processing pipeline. Pre-processing (Top, blue):** For each measurement, blip-up and blip-down diffusion volumes were corrected for eddy-current and subject motion using ECMOCO and subsequently combined into a susceptibility-distortion-corrected image with HySCO (using ACID toolbox in SPM12). The weighted average of the unwarped blip-up and blip-down volumes served as input to the DTI fit, yielding scalar maps (FA, MD, $\lambda_{1-3}$). **Brain-specific, longitudinal coregistration (Middle, green):** For every brain and time point ($t$: day 0/in-situ, day 0/beginning of fixation, ..., $n$/ days in fixative) the higher-resolution MTsat maps were first manually pre-aligned to the reference time point (in-situ for all brains except brain 5) and then refined by a contrast-based, threshold-driven automatic registration. The resulting transformation matrices were transferred to the corresponding b0/diffusion data, producing a longitudinally coregistered diffusion data set per brain (see Fritz et al. [38] for details). **Segmentation (Bottom, gray):** Mean FA and MD maps of the b=4000 s/mm$^2$ data were segmented into WM, cGM and dGM using SPM TPMs (95% probability threshold) in MNI space. The corresponding inverse deformation fields were applied to atlas labels (JHU for white-matter tracts, Harvard–Oxford for cortical and deep gray-matter regions) to project the ROIs (e.g. corpus callosum, putamen, thalamus) into specimen space. ROI boundaries were further constrained by an FA/MD-contrast-based expression on the mean maps and quality-controlled by visual inspection.

Figure S1 summarises the processing workflow used to obtain coregistered diffusion maps and anatomical ROIs for the longitudinal analyses. First, each ex-vivo multishell diffusion acquisition was split into single-shell datasets according to the analyzed b-values. Diffusion data were corrected for eddy-current and motion effects with ECMOCO and for susceptibility-related distortions with HySCO in the ACID toolbox. For the 2k4k acquisition, corrected blip-up and blip-down images were combined into a weighted-average image before tensor fitting. Because the 1k2k acquisition did not include blip-down images, the corrected 2k4k b0 images served as susceptibility-correction reference for the 1k2k blip-up data. DTI was then fitted with NLLS, yielding FA, MD, and eigenvalue maps.

Second, longitudinal coregistration was performed separately for each brain. The first available acquisition served as the reference time point, corresponding to the in-situ scan for all brains except brain 5. MPM acquisition and MTsat estimation followed Fritz et al. [38]. Because MTsat maps provide higher anatomical contrast and resolution than the diffusion data, they were first manually pre-aligned and then registered automatically across fixation time points. The resulting transformations were applied to the corresponding diffusion-derived maps, producing a coregistered temporal dataset for each brain.

Third, the coregistered diffusion maps were used for tissue segmentation and ROI definition. Brain-specific mean FA and MD maps were segmented into WM, cGM, and dGM using SPM tissue probability maps in MNI space (MNI) space with a 95 % probability threshold. Inverse deformation fields projected atlas labels from the JHU white-matter atlas and Harvard–Oxford gray-matter atlas into specimen space. The resulting ROIs, including the CC, internal capsule, and gray-matter regions, were refined using FA/MD contrast on the mean maps and visually quality-controlled.

##### 7.3.1 Used time intervals for analysis using tissue conditions

| | **in-situ 0 days** | **beginning of fixation 0 - 3 days** | **fixed 40-45 days** | **hydrated 1 - 42 days** |
|---|---|---|---|---|
| **brain 1:** | 1 dataset | 1 dataset | | 1 datasets |
| **brain 3:** | 1 dataset | 4 dataset | 4 datasets | 4 datasets |
| **brain 4:** | 1 dataset | | | |
| **brain 5:** | | 2 dataset | 2 datasets | 3 datasets |
| **brain 6** | 1 dataset | 2 dataset | 4 datasets | 3 datasets |

**Table S2:** Detailed information on which time intervals were used for Figure S2, as well as the number of data points per tissue condition and brain. Fixation was carried out with 4 % formaldehyde and hydrated with PBS.

#### 7.4 Protocol Comparison

The following sections contain the detailed methods and results of the comparative analysis of diffusion protocols. These analyses established that b-value 4000 s/mm$^2$ yielded the lowest RMSE and parameter variance across tissue conditions, which motivated its use for all main analyses in the manuscript.

##### 7.4.1 Methods: Comparative study of diffusion protocols

First, we compared diffusion protocols to assess how b-value affects diffusion parameter estimation and to identify the selected protocol for in-situ and early-fixation measurements. For this purpose, RMSE maps, reflecting the difference between measured and estimated signals, were compared across four protocols and tissue conditions: in-situ, beginning of fixation with formaldehyde (4–68 h after fixation onset), fixed (40–50 days in formaldehyde), and hydrated (around 9–27 days after transfer from formaldehyde to PBS).

Subsequently, voxelwise variation in MD and FA estimated with DTI was assessed across the same tissue conditions to evaluate the effects of lower-diffusion-weighting protocols.

Second, temporal evolution during fixation was modeled for all four diffusion protocols to examine how lower diffusion weighting affects the characterization of the fixation process. The main longitudinal and PMI-dependent analyses were then performed using the b-value 4000 s/mm$^2$ protocol. The following analyses all use the same saturation models (Section 2.5.2) and model selection procedure (Section 7.5.1).

**Protocol optimization: Examining the error of the DTI model for different diffusion protocols** The root mean square error (RMSE) is used to quantify the goodness of fit of the estimated diffusion maps for a given protocol. It is calculated with the error of the measured signal $\hat{S_n}^m$ and the estimated diffusion signal $S_n{}^e$ by using the DTI model as a sum over the diffusion directions $N$:

$$\mathrm{RMSE} = \sqrt{\frac{1}{N}\sum_{n=1}^{N}(\hat{S_n}^m - S_n{}^e)^2} \qquad \text{[S1]}$$

To compare between the diffusion protocols the TE-dependent b0-images were excluded for the RMSE calculation in Eq. S1, leading to a b-dependent RMSE. The lower the RMSE the better is the quality of the estimated DTI fit of the diffusion data, i.e., a large RMSE value is considered a poor estimate.

For the four aforementioned tissue conditions, mean b-dependent RMSE values were evaluated in WM to assess signal-fit quality across a broad white-matter region rather than a single tract. To assess statistical differences in the RMSE between imaging protocols, we employed a LME. The model was fitted with the corresponding protocol as a fixed effect and a random intercept for each brain to account for non-independence of repeated measurements from the same subject. Post-hoc pairwise comparisons between protocols were conducted using a coefficient test on the LME, with p-values adjusted for multiple comparisons using the Bonferroni correction.

**Influence of diffusion protocols on diffusion parameter estimates and variability** The diffusion parameters FA and MD were voxelwise evaluated over all brain samples in the CC ROI across the same four tissue conditions. In contrast to the WM-based RMSE analysis, parameter variance was assessed in the CC to minimize contributions from tissue heterogeneity. With the high alignment and homogeneous microstructural properties of the CC, observed variance in the data is more likely attributable to protocol-related model performance than underlying tissue heterogeneity. [39]

The variance of the data is graphically represented by boxplots for all four protocols. The whiskers' endpoints correspond to the lowest and highest values within 0.75 times the IQR (± 1.01 sigma, and 68.76 % coverage of the data). The lower quartile ($q_n(0.25)$) is the median of the lower half of the dataset, and the upper quartile ($q_n(0.75)$) is the median of the upper half. The IQR, representing the box's length, is the distance between ($q_n(0.25)$) and ($q_n(0.75)$) and contains the middle 50 % of the data points.

$$\mathrm{IQR} = q_{\mathrm{n}}(0.75) - q_{\mathrm{n}}(0.25) \qquad \text{[S2]}$$

##### 7.4.2 Supplementary Results: Comparative study of diffusion protocols

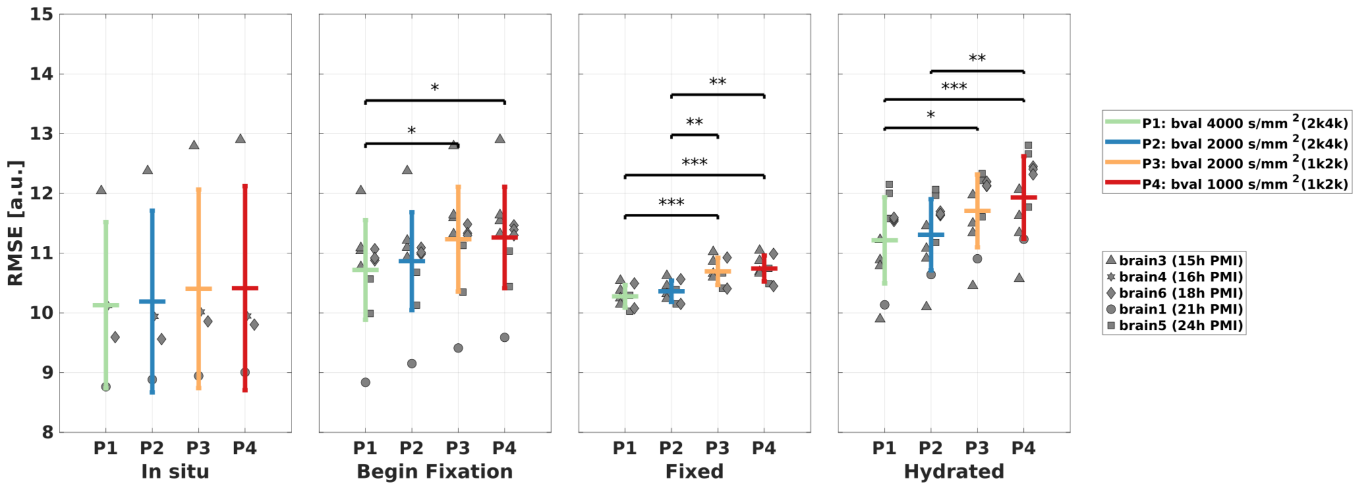


**Figure S2: Comparison of the mean RMSE values across diffusion protocols at different tissue conditions in the WM region:** Each subplot represents a distinct tissue condition: in-situ measurements, the beginning of fixation (0-3 days in 4% formaldehyde), fixed (40-45 days in 4 % formaldehyde), and hydrated in PBS.

The RMSE represents the error between the measured diffusion signal and the estimated diffusion signal from the DTI model. The colored error bars represent different protocols used for diffusion data acquisition: green (2k4k acquisition with b=4000 s/mm$^2$), blue (2k4k acquisition with b=2000 s/mm$^2$), yellow (1k2k acquisition with b=2000 s/mm$^2$), and red (1k2k acquisition with b=1000 s/mm$^2$). The horizontal lines indicate the mean RMSE across the WM of all brains for each protocol, while the vertical bars show the standard deviation.

Gray markers represent the underlying data from individual brains within the corresponding tissue condition (circle = brain1, triangle = brain3, star = brain4, square = brain5, diamond = brain6).

Statistical significance between protocols was determined using a LME Model with a Bonferroni correction for post-hoc pairwise comparisons. Significance levels are denoted as follows: * = statistically significant ($p < 0.05$), ** = highly significant ($p < 0.01$), *** = very highly significant ($p < 0.001$).

**Protocol optimization: Examining the error of the DTI model for different diffusion protocols** Figure S2 shows the mean root mean square error (RMSE) (see Eq.S1) across all brains for different tissue conditions and for four datasets derived from the two multishell acquisitions: b-value 4000 s/mm$^2$ (2k4k), b-value 2000 s/mm$^2$ (2k4k), b-value 2000 s/mm$^2$ (1k2k), and b-value 1000 s/mm$^2$ (1k2k).

The brain-wise averaged RMSE was lowest for b-value 4000 s/mm$^2$ and highest for b-value 1000 s/mm$^2$. Most informative for interpretation, the two datasets with identical nominal b-value 2000 s/mm$^2$ still differed: the 2k4k-derived shell was on average 3.28 % lower than the 1k2k-derived shell (2.13 % in-situ to 3.99 % hydrated). A linear mixed-effects model (LME) indicated no significant protocol effect in-situ ($p > 0.05$), but a significant effect at the other three tissue conditions.

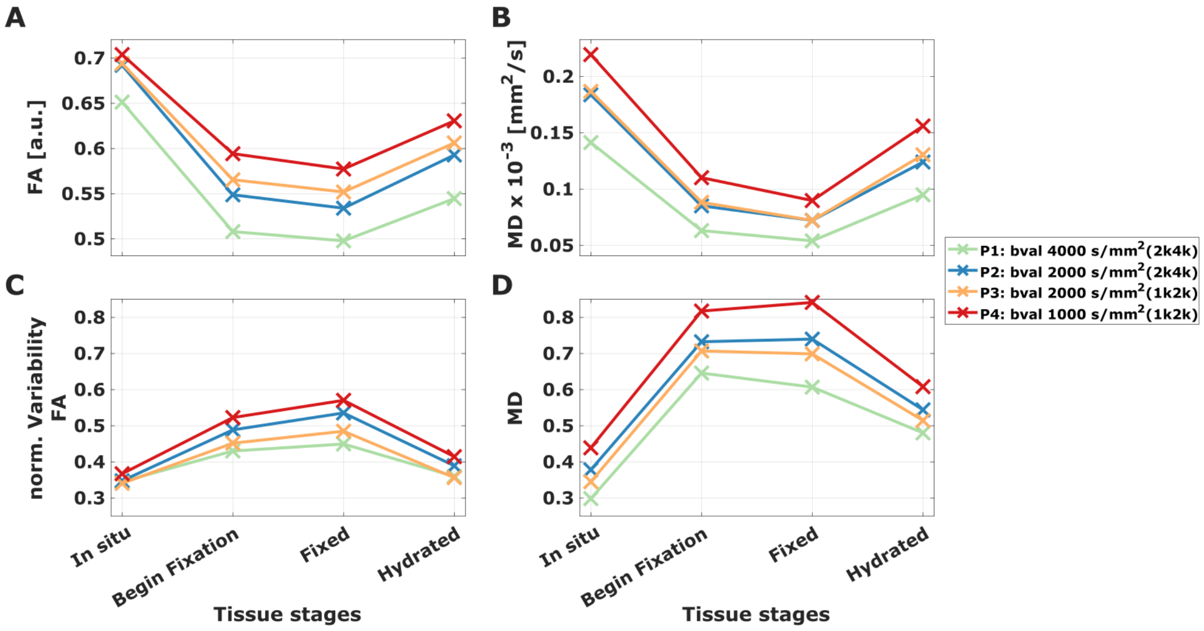


**Figure S3: Diffusion parameters and normalized variability in the CC:** Median values of FA (**A**) and MD (**B**) are shown in the top row, and the corresponding normalized IQR (normalized variability) of FA (**C**) and MD (**D**) in the bottom row. Values are plotted as line graphs for four single-shell datasets derived from the two multishell acquisitions across tissue conditions: in-situ measurements, beginning of fixation (0–3 days in 4% formaldehyde), fixed (40–45 days in 4% formaldehyde), and hydrated in PBS.

**Influence of diffusion protocols on diffusion parameter estimates and variability** For the median values of FA and MD (Figure S3 **A** and **B**), a consistent inverse relationship with the b-value was observed: the higher the b-value, the lower the estimated diffusion parameter across all tissue conditions. When comparing b-value 4000 s/mm$^2$ and b-value 1000 s/mm$^2$, the relative change in median FA reached a maximum decrease of 14.5 % at the beginning of fixation and a minimum decrease of 7.5 % in the in-situ condition. The two datasets with the same nominal b-value 2000 s/mm$^2$ (one from each multishell acquisition) yielded nearly identical median values across all tissue conditions. The relative change in median FA between these two b-value 2000 s/mm$^2$ datasets was minimal, ranging from $-0.27$ % (in-situ) to $-3.3$ % (fixed state). For MD, the two b-value 2000 s/mm$^2$ datasets also showed highly consistent results, ranging from $-4.9$ % (hydrated state) to 0.04 % (fixed state).

To assess variability in the diffusion parameters, we used the normalized IQR (IQR/median), which will be referred to as normalized variability in the following. As shown in Figure S3 **C** and **D**, both FA and MD exhibit a consistent trend across tissue conditions: the normalized variability increases from the in-situ to the fixed condition (on average across protocols by 97.6 % for MD and 45.7 % for FA) and decreases again from the fixed to the hydrated condition (by 25.6 % for MD and 25.5 % for FA). Among the two b-value 2000 s/mm$^2$ datasets, the 1k2k-derived dataset consistently showed lower normalized variability than the 2k4k-derived dataset across both diffusion parameters and all tissue conditions. Overall, b-value 4000 s/mm$^2$ exhibited the lowest normalized variability for both MD and FA compared to b-value 1000 s/mm$^2$, with differences for MD ranging from 21 % (beginning of fixation and hydrated) to 32 % (in-situ), and for FA most pronounced at the beginning of fixation (17.7 % lower) and in the fixed state (21.2 % lower).

The analysis of the IQR (Figure S3**C,D**) revealed that the lowest variability for both diffusion parameters across all tissue conditions was observed when the protocol with b-value 4000 s/mm$^2$ was used. Using a b-value much lower than 4000 s/mm$^2$ resulted in greater variance in the data.

The raw signal-fit RMSE is sensitive to the measured signal and is therefore expected to scale with b-value, because diffusion weighting changes signal attenuation and SNR. Although this b-value dependence is expected, the comparison of the two b-value 2000 s/mm$^2$ datasets indicates that the RMSE was more sensitive to the broader acquisition protocol than to nominal b-value alone: despite identical nominal diffusion weighting, their RMSE differed, indicating that residual protocol factors also affected signal-fit quality. In contrast, median FA and MD were nearly identical between these two b-value 2000 s/mm$^2$ datasets, suggesting that the absolute parameter offset was driven more strongly by b-value than by the remaining protocol differences. This protocol recommendation should be interpreted in the context of our acquisition setup and processing pipeline.

##### 7.4.3 Protocol comparison in corpus callosum and whole white matter

This section extends the CC-based b-value comparison shown in the main text (Figure 3) by including the second b-value 2000 s/mm$^2$ dataset, which originates from the 1k2k acquisition rather than the 2k4k acquisition used in the main analyses. By plotting both b-value 2000 s/mm$^2$ datasets side by side at otherwise identical fixed nominal diffusion weighting, the comparison isolates the effect of remaining protocol differences (TE/TR, distortion-correction conditions, blip-up/-down availability) from the effect of the b-value itself. Figures S5 and S4 show the CC analysis for FA and MD, respectively, and Figures S7 and S6 report the analogous comparison in whole-brain WM. Each figure shows all four analyzed datasets (b-value 4000 and 2000 s/mm$^2$ from the 2k4k acquisition and b-value 2000 and 1000 s/mm$^2$ from the 1k2k acquisition) as panels **A**–**D**, together with the fixation-model nRMSE as a function of b-value (panel **E**).

**Corpus callosum: fixation sensitivity and precision across b-values.** All four datasets captured the fixation effect in the CC: MD decreased sharply toward a plateau (Figure S4) and FA decreased or stabilized after an initial transient (Figure S5), confirming that sensitivity to fixation is not restricted to a particular diffusion weighting. Precision, however, increased markedly with b-value: lower b-values produced a systematic positive value offset and larger scatter across specimens and adjacent time points in both MD and FA, whereas higher b-values yielded tighter trajectories. For MD, the fixation-model nRMSE decreased monotonically with b-value, being highest at b-value 1000 s/mm$^2$ and lowest at b-value 4000 s/mm$^2$ (Figure S4**E**). For FA, the nRMSE remained comparatively high across all b-values without consistent improvement (Figure S5**E**).

**Protocol-dependence at fixed b-value in the CC.** For both FA and MD, the time courses for the two b-value 2000 s/mm$^2$ datasets were qualitatively similar but not identical, and a systematic offset in the fixation-model nRMSE was observed (Figure S5**E** and Figure S4**E**). The b-value 2000 s/mm$^2$ shell derived from the 2k4k acquisition yielded a lower nRMSE in MD than the b-value 2000 s/mm$^2$ shell derived from the 1k2k acquisition, despite sharing the same nominal diffusion weighting. This indicates that, in addition to the dominant b-value effect, residual protocol-related factors (TE/TR, distortion correction with/without blip-down reference) contribute to the apparent fit quality. The b-value comparison shown in the main text (Figure 3) therefore captures the principal protocol effect, but a smaller, protocol-specific contribution exists on top of it and is recoverable here.

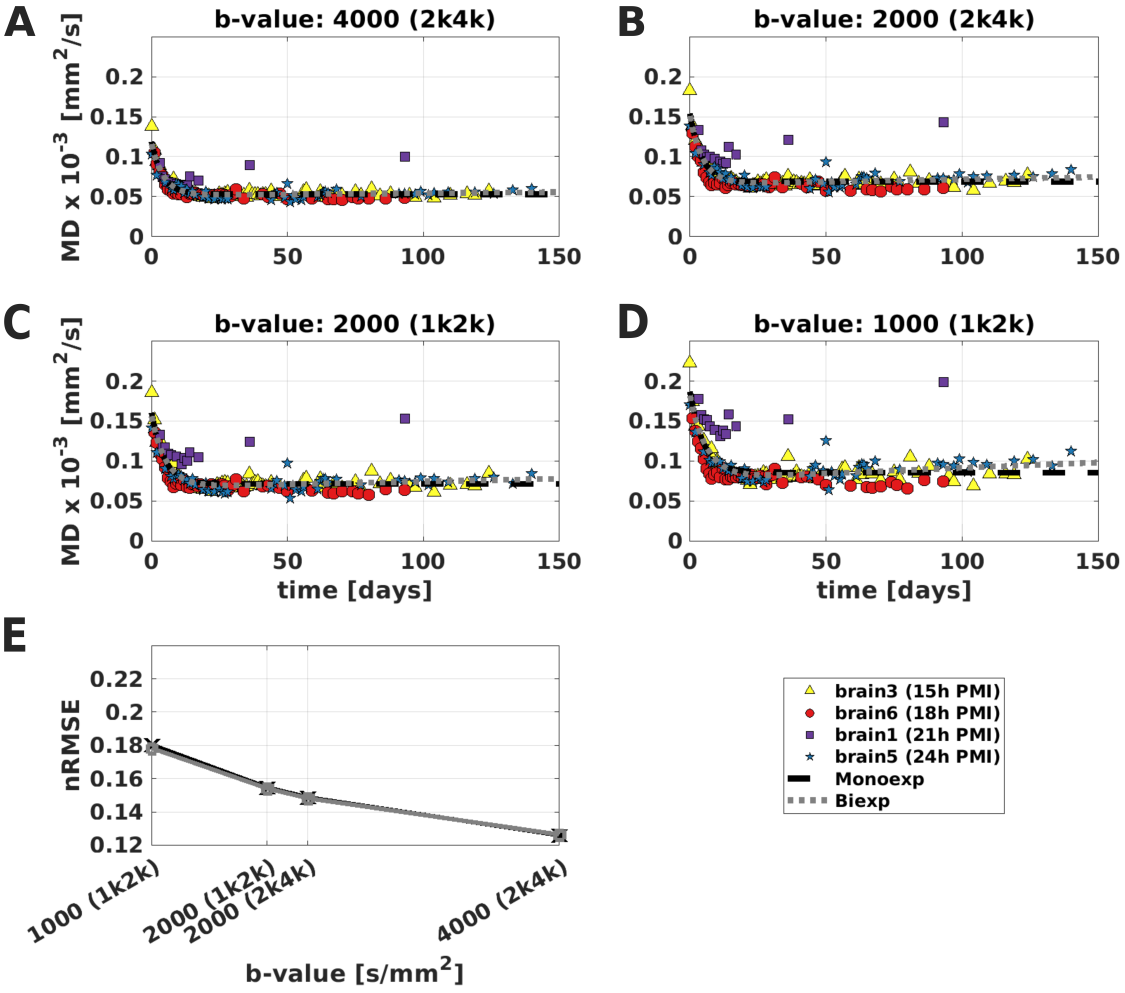


**Figure S4: Temporal evolution modeling of mean diffusivity (MD) during fixation in the corpus callosum (CC): comparison across all four multishell datasets (Supplement to Figure 3).** (**A**)–(**D**) Median MD over fixation time (days) for (**A**) b-value 4000 s/mm$^2$ (2k4k), (**B**) b-value 2000 s/mm$^2$ (2k4k), (**C**) b-value 2000 s/mm$^2$ (1k2k), and (**D**) b-value 1000 s/mm$^2$ (1k2k). Data from four brains with different post-mortem intervals (PMI): brain3 (15 h, yellow triangles), brain6 (18 h, red circles), brain1 (21 h, purple squares), and brain5 (24 h, blue stars). Black solid line: monoexponential fit; gray dotted line: biexponential fit. (**E**) Normalized root mean square error (nRMSE) of the fixation model as a function of b-value, comparing monoexponential (black, crosses) and biexponential (gray, squares) fits; the two points at b-value 2000 s/mm$^2$ originate from the 2k4k and 1k2k acquisitions and illustrate the residual protocol-dependence at matched nominal diffusion weighting. nRMSE was computed as in Supporting Information (Section 2.5.2 of the Methods).

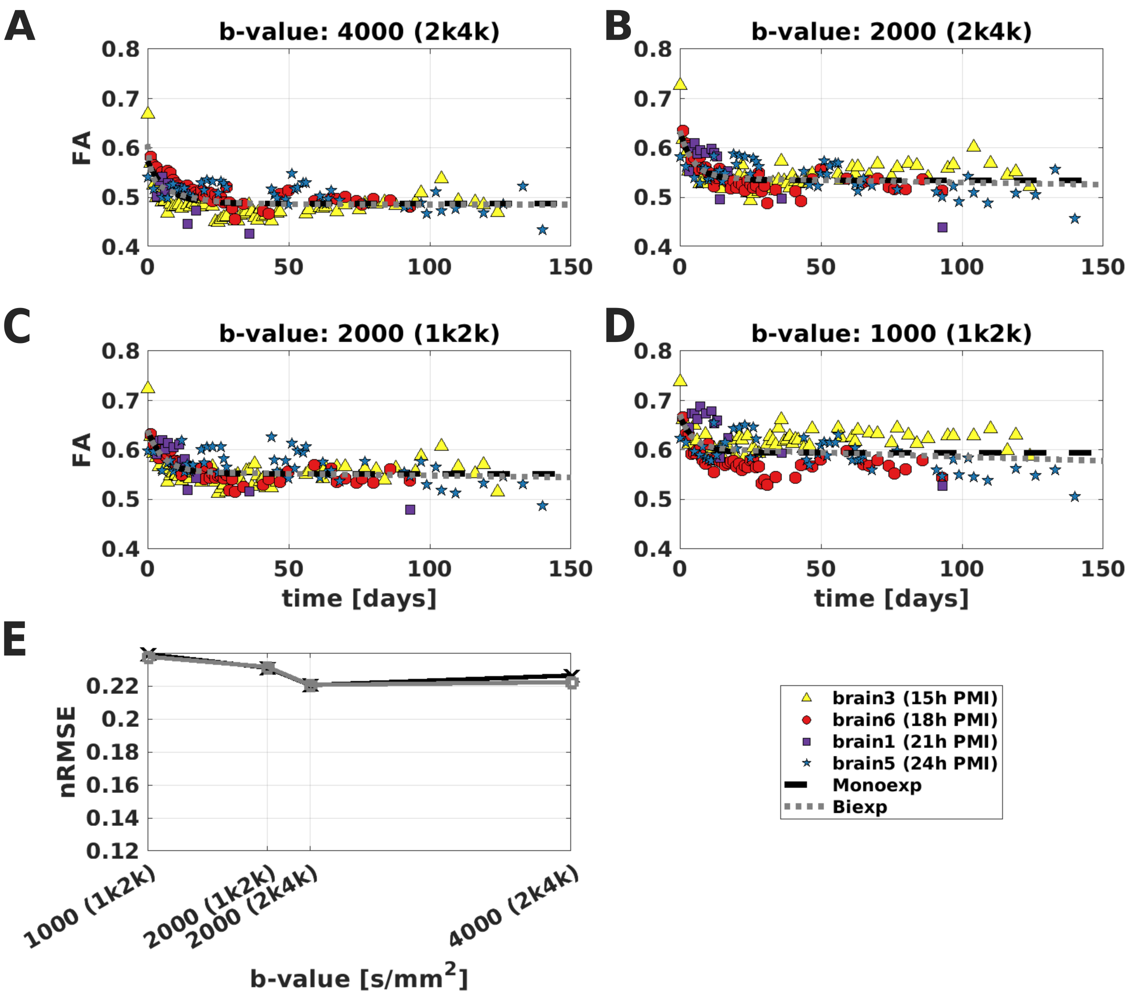


**Figure S5: Temporal evolution modeling of fractional anisotropy (FA) during fixation in the corpus callosum (CC): comparison across all four multishell datasets (Supplement to Figure 3).** (**A**)–(**D**) Median FA over fixation time (days) for (**A**) b-value 4000 s/mm$^2$ (2k4k), (**B**) b-value 2000 s/mm$^2$ (2k4k), (**C**) b-value 2000 s/mm$^2$ (1k2k), and (**D**) b-value 1000 s/mm$^2$ (1k2k). Data from four brains with different post-mortem intervals (PMI): brain3 (15 h, yellow triangles), brain6 (18 h, red circles), brain1 (21 h, purple squares), and brain5 (24 h, blue stars). Black solid line: monoexponential fit; gray dotted line: biexponential fit. (**E**) Normalized root mean square error (nRMSE) of the fixation model as a function of b-value, comparing monoexponential (black, crosses) and biexponential (gray, squares) fits; the two points at b-value 2000 s/mm$^2$ originate from the 2k4k and 1k2k acquisitions and illustrate the residual protocol-dependence at matched nominal diffusion weighting. nRMSE was computed as in Supporting Information (Section 2.5.2 of the Methods).

**Whole-brain WM: same trend, larger heterogeneity.** Figures S7 and S6 show the temporal evolution of FA and MD during fixation in whole-brain WM, again including all four datasets and both b-value 2000 s/mm$^2$ shells in the nRMSE panel.

For MD, WM followed the same exponential-like decrease toward a plateau as the CC, although the decline was somewhat slower and reached a slightly higher plateau (Figure S6**A**–**D**), with monoexponential and biexponential fits remaining nearly indistinguishable. As in the CC, lower b-values produced a positive value offset and larger scatter, and the nRMSE for MD decreased with increasing b-value (Figure S6**E**); the protocol-dependence between the two b-value 2000 s/mm$^2$ datasets reproduced the offset observed in the CC.

For FA, the between-brain heterogeneity was even more pronounced in WM than in the CC. Importantly, the two specimens with longer PMI (brain1, 21 h, and brain5, 24 h) showed an upward drift in FA from the beginning of fixation across all datapoints, whereas the two specimens with shorter PMI (brain3, 15 h, and brain6, 18 h) showed an initial decrease over the first days before stabilizing. This PMI-dependent divergence in FA direction was visible at all b-values but was most pronounced at b-value 1000 s/mm$^2$, where the overall FA level was additionally offset upward and the larger inter-specimen scatter makes individual trends harder to separate from noise (Figure S7**A**–**D**). As a consequence, no single saturation model described all brains simultaneously, and the nRMSE for FA remained comparatively high and showed no consistent improvement with increasing b-value (Figure S7**E**). This non-uniform FA behavior in WM motivated the subsequent stratification of WM by fiber orientation dispersion ($\kappa$) in the main analyses (Section 3.6).

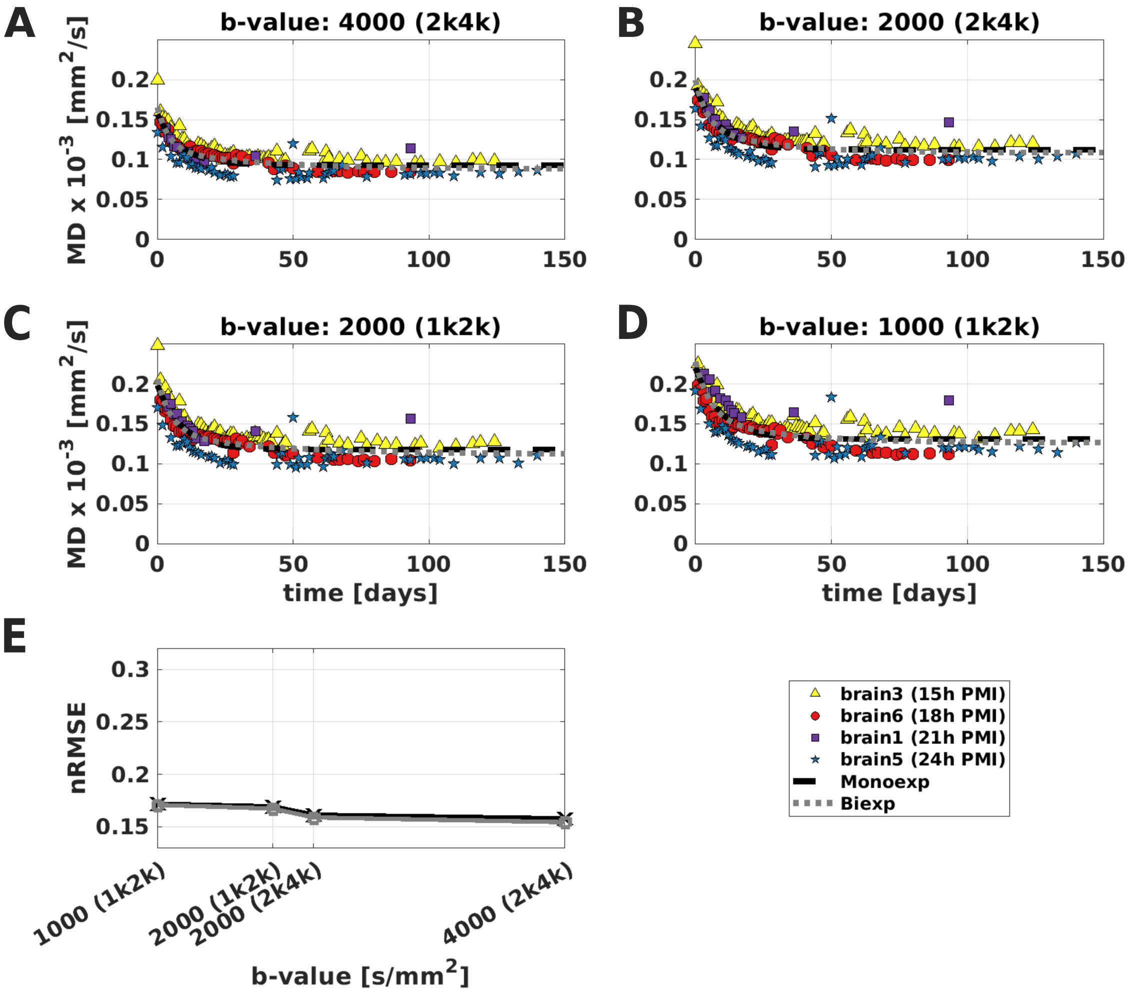


**Figure S6: Temporal evolution modeling of mean diffusivity (MD) during fixation in whole-brain white matter (WM): comparison across all four multishell datasets (Supplement to Figure 3).** (**A**)–(**D**) Median MD over fixation time (days) for (**A**) b-value 4000 s/mm$^2$ (2k4k), (**B**) b-value 2000 s/mm$^2$ (2k4k), (**C**) b-value 2000 s/mm$^2$ (1k2k), and (**D**) b-value 1000 s/mm$^2$ (1k2k). Data from four brains with different post-mortem intervals (PMI): brain3 (15 h, yellow triangles), brain6 (18 h, red circles), brain1 (21 h, purple squares), and brain5 (24 h, blue stars). Black solid line: monoexponential fit; gray dotted line: biexponential fit. (**E**) Normalized root mean square error (nRMSE) of the fixation model as a function of b-value, comparing monoexponential (black, crosses) and biexponential (gray, squares) fits; the two points at b-value 2000 s/mm$^2$ originate from the 2k4k and 1k2k acquisitions. nRMSE was computed as in Supporting Information (Section 2.5.2 of the Methods).

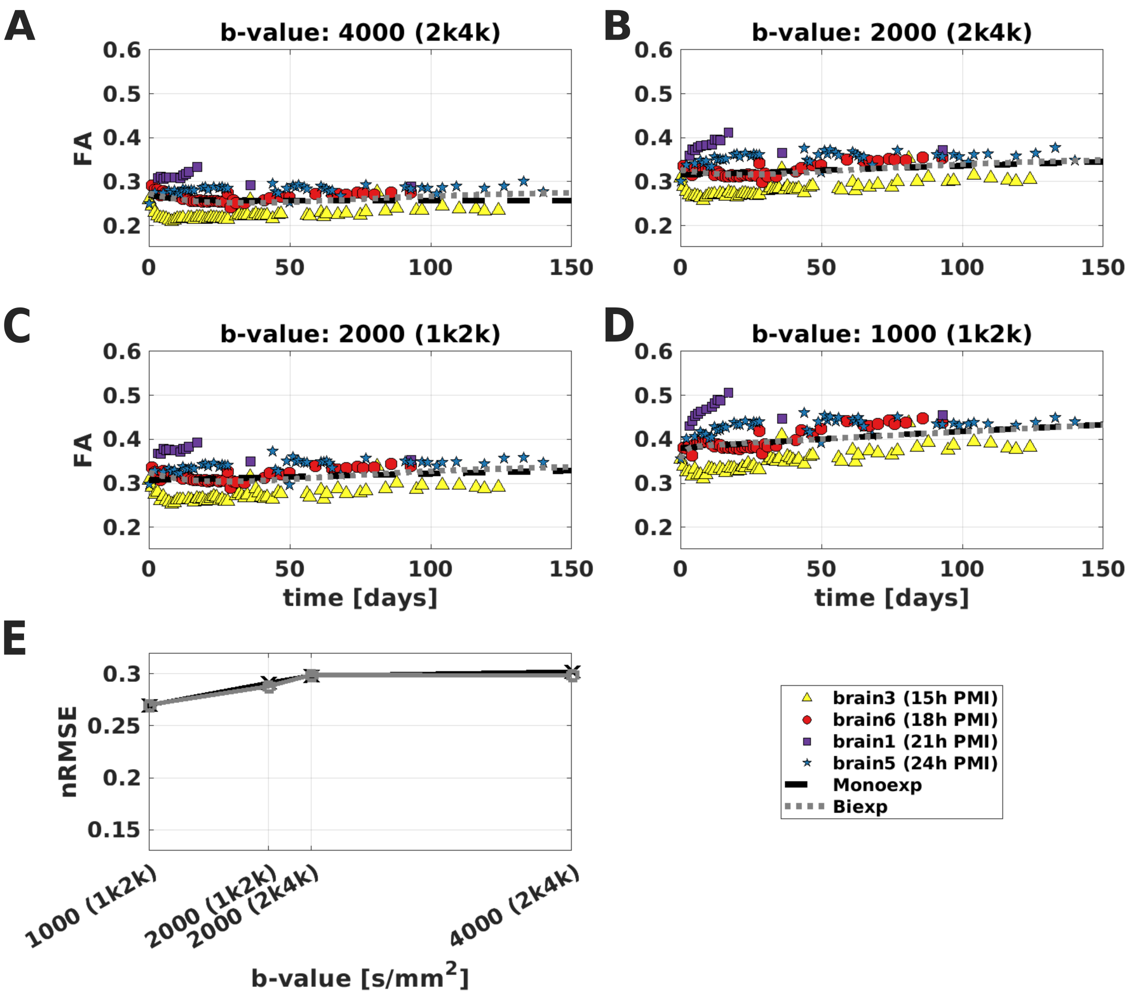


**Figure S7: Temporal evolution modeling of fractional anisotropy (FA) during fixation in whole-brain white matter (WM): comparison across all four multishell datasets (Supplement to Figure 3).** (**A**)–(**D**) Median FA over fixation time (days) for (**A**) b-value 4000 s/mm$^2$ (2k4k), (**B**) b-value 2000 s/mm$^2$ (2k4k), (**C**) b-value 2000 s/mm$^2$ (1k2k), and (**D**) b-value 1000 s/mm$^2$ (1k2k). Data from four brains with different post-mortem intervals (PMI): brain3 (15 h, yellow triangles), brain6 (18 h, red circles), brain1 (21 h, purple squares), and brain5 (24 h, blue stars). Black solid line: monoexponential fit; gray dotted line: biexponential fit. (**E**) Normalized root mean square error (nRMSE) of the fixation model as a function of b-value, comparing monoexponential (black, crosses) and biexponential (gray, squares) fits; the two points at b-value 2000 s/mm$^2$ originate from the 2k4k and 1k2k acquisitions. nRMSE was computed as in Supporting Information (Section 2.5.2 of the Methods).

Although both MD and FA are derived from the same diffusion-tensor eigenvalues ($\lambda_1, \lambda_2, \lambda_3$), their fixation-time behavior and model-fit characteristics differed. In the eigenvalue analysis, all three eigenvalues decreased during fixation, see Appendix 6.2. However, this common directional trend translated into a comparatively stable and well-modeled MD trajectory, while FA remained more heterogeneous across brains and regions. This difference is expected because MD reflects the average magnitude of diffusion, whereas FA depends on the relative spacing of the eigenvalues and is therefore more sensitive to subtle differences in anisotropy and tissue organization.

A further observation is that the protocol-related nRMSE reduction for MD was more pronounced in the CC than in whole-brain WM. A likely explanation is the different number of voxels entering the ROI summary: whole-WM contains substantially more voxels and therefore benefits from stronger averaging and a higher effective SNR, which stabilises estimates already at lower diffusion weighting. In the CC, where fewer voxels contribute to each time-point summary, precision is more sensitive to protocol choice. Consequently, the gain from ex-vivo-adjusted diffusion weighting is more visible in nRMSE. In practical terms, this suggests that protocol optimization can provide a larger precision benefit in smaller ROIs, whereas the relative gain is attenuated in very large ROIs due to voxel averaging.

#### 7.5 Modeling the temporal evolution of diffusion parameters during fixation and protocol comparison

##### 7.5.1 Model selection and evidence ratios

**Methods:** To model the temporal evolution of FA and MD during fixation, model preference was assessed using an information-theoretic approach. The candidate models were the null model, the monoexponential saturation model, and the biexponential saturation model defined in the Methods (Section 2.5.2). For each ROI, diffusion parameter, and acquisition protocol, models were fitted to the longitudinal diffusion data during immersion fixation across specimens. Model preference was first quantified by the Akaike Information Criterion (AIC), then corrected for finite sample size using the corrected Akaike Information Criterion (AICc), and finally expressed as evidence ratios (ER) relative to the best-supported model. [47, 48]

The AIC for model $m$ was computed as

$$\mathrm{AIC}_m = n\ln(\mathrm{RSS}_m/n) + 2k_m, \qquad \text{[S3]}$$

where $k_m$ is the number of fitted parameters, $n$ is the number of observations, and $\mathrm{RSS}_m$ is the residual sum of squares between the fitted model and the measured data. Lower AIC values indicate a better trade-off between goodness of fit and model complexity.

Because the number of observations was finite, AIC was corrected to AICc: [49, 50]

$$\mathrm{AICc}_m = \mathrm{AIC}_m + \frac{2k_m(k_m+1)}{n-k_m-1}. \qquad \text{[S4]}$$

The model with the lowest AICc was considered the best-supported model by this criterion. [51]

To express the strength of evidence against each non-best model, evidence ratios were calculated from the AICc differences: [47, 48, 49]

$$\mathrm{ER}_m = \exp\left(\frac{\mathrm{AICc}_m - \mathrm{AICc}_{\mathrm{min}}}{2}\right), \qquad \text{[S5]}$$

where $\mathrm{AICc}_{\mathrm{min}}$ is the lowest AICc across the candidate models. The lowest-AICc model therefore has $\mathrm{ER} = 1$. Models with $\mathrm{ER} < 2.71$ were considered to have substantial support relative to the lowest-AICc model, whereas models with $\mathrm{ER} > 148.41$ were considered to have essentially no support. This model-selection framework was used to determine which model is shown in the main text, whereas the Supporting Information Figures below show monoexponential and biexponential alternatives side by side for transparency. The resulting AICc and ER values are summarized in Table S3.

| | | | | AICc | | | ER | | | |
|---|---|---|---|---|---|---|---|---|---|---|
| ROI | Parameter | Protocol | *n* | Null | Mono | Biexp | Null | Mono | Biexp | Preferred |
| Corpus callosum (CC) | FA | 4000 (2k4k) | 169 | -1170.08 | -1237.68 | **-1241.56** | $\gg 10^4$ | 6.98 | **1.00** | Biexp |
| | | 2000 (2k4k) | 169 | -1157.72 | **-1227.09** | -1224.77 | $\gg 10^4$ | **1.00** | 3.18 | Mono |
| | | 2000 (1k2k) | 169 | -1177.87 | **-1245.81** | -1242.93 | $\gg 10^4$ | **1.00** | 4.22 | Mono |
| | | 1000 (1k2k) | 169 | -1123.99 | **-1151.63** | -1151.61 | $\gg 10^4$ | **1.00** | 1.01 | Mono |
| | MD | 4000 (2k4k) | 169 | -1454.99 | **-1679.50** | -1676.30 | $\gg 10^4$ | **1.00** | 4.95 | Mono |
| | | 2000 (2k4k) | 169 | -1336.63 | **-1505.98** | -1504.99 | $\gg 10^4$ | **1.00** | 1.64 | Mono |
| | | 2000 (1k2k) | 169 | -1313.34 | **-1489.20** | -1488.28 | $\gg 10^4$ | **1.00** | 1.58 | Mono |
| | | 1000 (1k2k) | 169 | -1222.22 | -1344.78 | **-1346.41** | $\gg 10^4$ | 2.27 | **1.00** | Mono |
| White matter (WM) | FA | 4000 (2k4k) | 169 | **-1184.29** | -1181.94 | -1183.73 | **1.00** | 3.24 | 1.32 | Null |
| | | 2000 (2k4k) | 169 | -1115.28 | **-1117.24** | -1115.38 | 2.66 | **1.00** | 2.53 | Null |
| | | 2000 (1k2k) | 169 | -1132.18 | -1131.74 | **-1133.47** | 1.91 | 2.37 | **1.00** | Null |
| | | 1000 (1k2k) | 169 | -1055.14 | **-1065.58** | -1062.86 | 185 | **1.00** | 3.91 | Mono |
| | MD | 4000 (2k4k) | 169 | -1341.41 | -1506.68 | **-1512.47** | $\gg 10^4$ | 18.0 | **1.00** | Biexp |
| | | 2000 (2k4k) | 169 | -1273.70 | -1443.08 | **-1446.34** | $\gg 10^4$ | 5.10 | **1.00** | Biexp |
| | | 2000 (1k2k) | 169 | -1258.69 | -1421.48 | **-1423.65** | $\gg 10^4$ | 2.96 | **1.00** | Biexp |
| | | 1000 (1k2k) | 169 | -1214.70 | **-1376.26** | -1376.14 | $\gg 10^4$ | **1.00** | 1.06 | Mono |

**Table S3:** Summary of the corrected Akaike Information Criterion (AICc) and evidence ratios (ER) for null, monoexponential, and biexponential models fitted to the temporal change in FA and MD during fixation in corpus callosum (CC) and white matter (WM) across DTI acquisition protocols. Lower (more negative) AICc indicates better fit. **Bold AICc** marks the lowest AICc per row. **Bold ER** (= 1.00) marks the lowest-AICc reference model for the evidence ratio. ER $< 2.71$ indicates substantial support relative to the lowest-AICc model; ER $> 148.41$ indicates essentially no support. Protocol notation: shell (gradient scheme), e.g. 4000 (2k4k) = $b = 4000$ s/mm$^2$ with 2k4k gradient scheme.

**Results:** In the CC, MD was best summarized by the monoexponential model, even when the biexponential model had a marginally lower AICc at b-value 1000 s/mm$^2$. For FA in the CC, model preference was less uniform: the biexponential model was preferred at b-value 4000 s/mm$^2$, whereas monoexponential fits were preferred at lower b-values. In whole-brain WM, FA often favored the null or simpler models, consistent with the heterogeneous and non-uniform time courses described below. In contrast, whole-brain WM MD retained a clear fixation-time dependence, with biexponential fits preferred for the higher b-values and a monoexponential fit preferred at b-value 1000 s/mm$^2$.

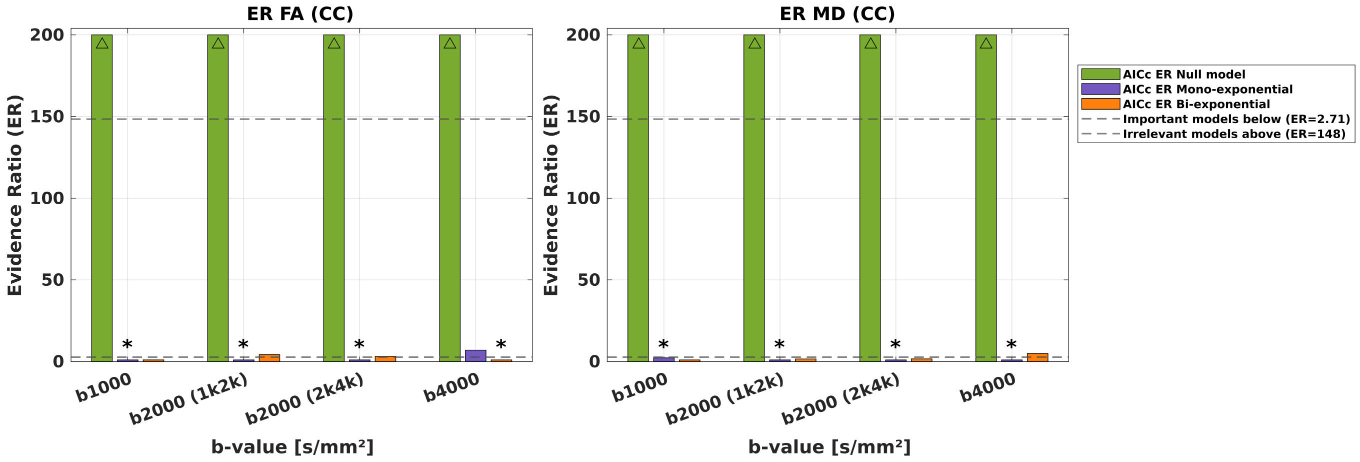


**Figure S8: Evidence ratios for fixation-model selection in the corpus callosum (CC).** Evidence ratios are shown for null, monoexponential, and biexponential models fitted to fixation-related fractional anisotropy (FA) and mean diffusivity (MD) changes in the CC across the analyzed diffusion protocols. ER values were computed from AICc differences relative to the lowest-AICc model (ER = 1). Dashed reference lines indicate ER = 2.71, below which models retain substantial support, and ER = 148.41, above which models have essentially no support relative to the lowest-AICc model.

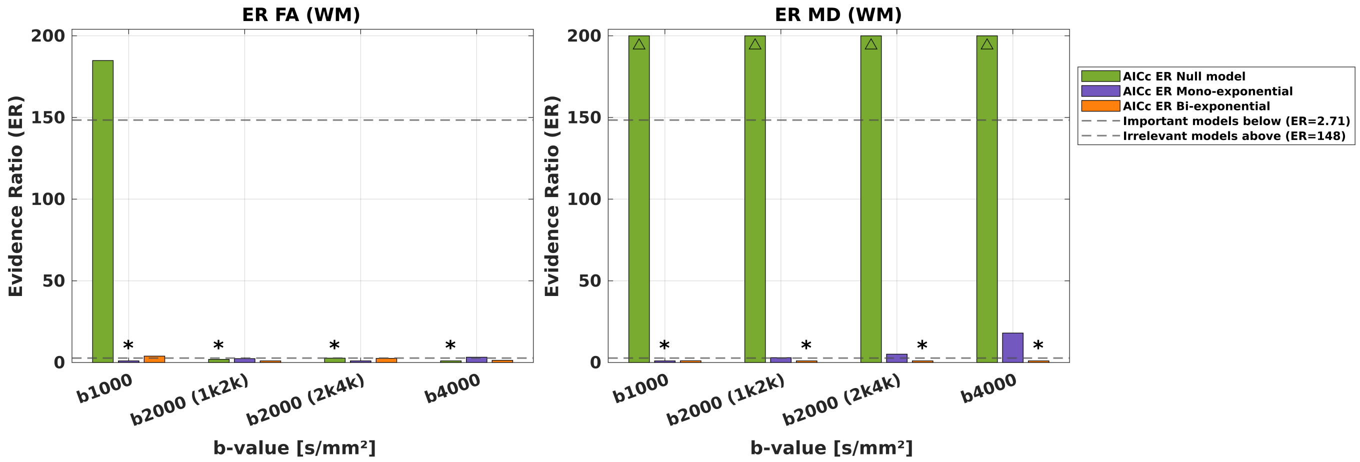


**Figure S9: Evidence ratios for fixation-model selection in whole-brain white matter (WM).** Evidence ratios are shown for null, monoexponential, and biexponential models fitted to fixation-related fractional anisotropy (FA) and mean diffusivity (MD) changes in whole-brain WM across the analyzed diffusion protocols. ER values were computed from AICc differences relative to the lowest-AICc model (ER = 1). Dashed reference lines indicate ER = 2.71, below which models retain substantial support, and ER = 148.41, above which models have essentially no support relative to the lowest-AICc model.

To make the preferred curves reproducible and to document cases in which the biexponential model introduced slow components with limited additional support, Table S4 reports fitted coefficients for both candidate saturation models.

| ROI | Parameter | Protocol | Model | $a$ | $b$ | $c$ | $d$ | $e$ |
|---|---|---|---|---|---|---|---|---|
| Corpus callosum (CC) | FA | 4000 (2k4k) | Mono | 0.5735 | 0.0857 | 0.1496 | — | — |
| | | | **Biexp*** | **0.6081** | **0.0687** | **0.9399** | **0.0535** | **0.0847** |
| | | 2000 (2k4k) | **Mono*** | **0.6348** | **0.0998** | **0.1981** | — | — |
| | | | Biexp | 0.6348 | 0.0948 | 0.2170 | 0.0991 | 0.0010 |
| | | 2000 (1k2k) | **Mono*** | **0.6404** | **0.0885** | **0.1727** | — | — |
| | | | Biexp | 0.6398 | 0.0838 | 0.1850 | 0.0789 | 0.0010 |
| | | 1000 (1k2k) | **Mono*** | **0.6687** | **0.0744** | **0.1818** | — | — |
| | | | Biexp | 0.6721 | 0.0684 | 0.2418 | 0.1829 | 0.0010 |
| | MD | 4000 (2k4k) | **Mono*** | **0.1188** | **0.0658** | **0.2509** | — | — |
| | | | Biexp | 0.1158 | 0.0646 | 0.2204 | -0.0290 | 0.0011 |
| | | 2000 (2k4k) | **Mono*** | **0.1540** | **0.0852** | **0.2386** | — | — |
| | | | Biexp | 0.1510 | 0.0857 | 0.2084 | -0.0550 | 0.0012 |
| | | 2000 (1k2k) | **Mono*** | **0.1605** | **0.0893** | **0.2197** | — | — |
| | | | Biexp | 0.1562 | 0.0895 | 0.1848 | -0.0796 | 0.0010 |
| | | 1000 (1k2k) | **Mono*** | **0.1869** | **0.1015** | **0.2014** | — | — |
| | | | Biexp | 0.1819 | 0.1050 | 0.1609 | -0.1504 | 0.0010 |
| White matter (WM) | FA | 4000 (2k4k) | Mono | 0.2700 | 0.0131 | 0.1159 | — | — |
| | | | Biexp | 0.2736 | 0.1711 | 0.0332 | -0.1777 | 0.0221 |
| | | 2000 (2k4k) | Mono | 0.3154 | -0.2167 | 0.0010 | — | — |
| | | | Biexp | 0.3270 | 0.2213 | 0.0313 | -0.2487 | 0.0232 |
| | | 2000 (1k2k) | Mono | 0.3068 | -0.1553 | 0.0010 | — | — |
| | | | Biexp | 0.3245 | 0.2737 | 0.0337 | -0.2906 | 0.0251 |
| | | 1000 (1k2k) | **Mono*** | **0.3799** | **-0.1491** | **0.0029** | — | — |
| | | | Biexp | 0.3545 | -0.0284 | 0.5754 | -0.3589 | 0.0010 |
| | MD | 4000 (2k4k) | Mono | 0.1569 | 0.0639 | 0.1157 | — | — |
| | | | **Biexp*** | **0.1654** | **0.0533** | **0.2299** | **0.0240** | **0.0289** |
| | | 2000 (2k4k) | Mono | 0.1898 | 0.0774 | 0.1104 | — | — |
| | | | **Biexp*** | **0.1999** | **0.0589** | **0.2359** | **0.0326** | **0.0390** |
| | | 2000 (1k2k) | Mono | 0.1985 | 0.0805 | 0.1124 | — | — |
| | | | **Biexp*** | **0.2060** | **0.0681** | **0.1902** | **0.0259** | **0.0257** |
| | | 1000 (1k2k) | **Mono*** | **0.2202** | **0.0894** | **0.1046** | — | — |
| | | | Biexp | 0.2279 | 0.0700 | 0.1819 | 0.0315 | 0.0343 |

**Table S4:** Estimated coefficients of the monoexponential and biexponential models fitted to the temporal change of FA and MD during fixation in corpus callosum (CC) and white matter (WM). Monoexponential model: $y = a - b\big(1 - \mathrm{e}^{-cx}\big)$. Biexponential model: $y = a - b\big(1 - \mathrm{e}^{-cx}\big) - d\big(1 - \mathrm{e}^{-ex}\big)$, with $x$ in days. **Bold** values (and *) mark the preferred model according to AICc with the parsimony rule ($\Delta$AICc $\leq 2$). For monoexponential fits, $d$ and $e$ are not applicable (—). Protocol notation: shell (gradient scheme), e.g. 4000 (2k4k) = $b = 4000$ s/mm$^2$ with 2k4k gradient scheme. MD values are scaled by $10^3$ (units: $10^{-3}$ mm$^2$/s); FA is dimensionless; rate constants $c$ and $e$ are in day$^{-1}$.

#### 7.6 Supplementary statistics: PMI associations with fixation-model parameters

Figure 4 relates brain-specific FA fixation-model parameters in the CC to PMI. Brain-specific saturation models were fitted at b-value 4000 s/mm$^2$ (2k4k protocol) using the same null, monoexponential, and biexponential candidates and AICc/ER framework as in Section 7.5.1. Model preference between monoexponential and biexponential alternatives followed the same parsimony rule ($\Delta$AICc $\leq 2$). Table S5 and Table S6 report the resulting AICc/ER values and fitted coefficients for brain3 (15 h), brain6 (18 h), brain1 (21 h), and brain5 (24 h PMI). Brain2 was excluded because it contributed too few fixation time points for reliable model fitting. Table S7 lists the OLS regression and Pearson correlation statistics across these four brains. The OLS regression line can be read as $Y = m\mathrm{PMI} + c$, where $Y$ is the fitted fixation parameter, $m$ is the slope, and $c$ is the intercept.

| | | | | AICc | | | ER | | | |
|---|---|---|---|---|---|---|---|---|---|---|
| ROI | Parameter | Brain (PMI) | *n* | Null | Mono | Biexp | Null | Mono | Biexp | Preferred |
| Corpus callosum (CC) | FA | brain3 (15 h) | 57 | -391.74 | **-481.21** | -478.14 | $\gg 10^4$ | **1.00** | 4.63 | Mono |
| | | brain6 (18 h) | 45 | -324.75 | **-396.55** | -394.04 | $\gg 10^4$ | **1.00** | 3.50 | Mono |
| | | brain1 (21 h) | 13 | -78.14 | **-95.51** | -90.72 | 5915 | **1.00** | 10.93 | Mono |
| | | brain5 (24 h) | 42 | -334.90 | -345.39 | **-348.47** | 886 | 4.67 | **1.00** | Biexp |

**Table S5:** Summary of the corrected Akaike Information Criterion (AICc) and evidence ratios (ER) for null, monoexponential, and biexponential models fitted to brain-specific FA time courses during fixation in the corpus callosum (CC) at b-value 4000 s/mm$^2$ (2k4k protocol). Lower (more negative) AICc indicates better fit. **Bold AICc** marks the lowest AICc per row. **Bold ER** (= 1.00) marks the lowest-AICc reference model for the evidence ratio. ER $< 2.71$ indicates substantial support relative to the lowest-AICc model; ER $> 148.41$ indicates essentially no support. The *Preferred* column reports the monoexponential/biexponential choice after the parsimony rule ($\Delta$AICc $\leq 2$).

| ROI | Parameter | Brain (PMI) | Model | *a* | *b* | *c* | *d* | *e* |
|---|---|---|---|---|---|---|---|---|
| Corpus callosum (CC) | FA | brain3 (15 h) | **Mono**$^*$ | **0.6525** | **0.1769** | **0.5000** | — | — |
| | | | Biexp | 0.6548 | 0.1836 | 0.5000 | -0.0150 | 0.0100 |
| | | brain6 (18 h) | **Mono**$^*$ | **0.5865** | **0.0967** | **0.0923** | — | — |
| | | | Biexp | 0.5822 | 0.3567 | 0.0489 | -0.2751 | 0.0373 |
| | | brain1 (21 h) | **Mono**$^*$ | **0.5502** | **0.1812** | **0.0291** | — | — |
| | | | Biexp | 0.3080 | -0.3027 | 0.5000 | 0.2240 | 0.0561 |
| | | brain5 (24 h) | Mono | 0.5265 | 0.0545 | 0.0100 | — | — |
| | | | **Biexp**$^*$ | **0.5162** | **-0.2712** | **0.0167** | **0.4055** | **0.0100** |

**Table S6:** Estimated coefficients of the monoexponential and biexponential models fitted to brain-specific FA time courses during fixation in the corpus callosum (CC) at b-value 4000 s/mm$^2$ (2k4k protocol). Monoexponential model: $y = a - b\left(1 - \mathrm{e}^{-cx}\right)$. Biexponential model: $y = a - b\left(1 - \mathrm{e}^{-cx}\right) - d\left(1 - \mathrm{e}^{-ex}\right)$, with $x$ in days. **Bold** values (and $^*$) mark the preferred monoexponential or biexponential model according to AICc with the parsimony rule ($\Delta$AICc $\leq 2$). For monoexponential fits, $d$ and $e$ are not applicable (—). FA is dimensionless; rate constants $c$ and $e$ are in day$^{-1}$.

| Region | Parameter | Symbol | $n$ | OLS $m$ | OLS $c$ | OLS $p$ | OLS $R^2$ | Pearson $r$ | Pearson $p$ |
|---|---|---|---|---|---|---|---|---|---|
| CC | Initial FA | FA($t_0$) | 4 | -0.014837 | 0.8657 | **0.014** | 0.971 | -0.986 | **0.014** |
| CC | Amplitude | ΔFA | 4 | -0.0014376 | 0.1753 | 0.860 | 0.020 | -0.140 | 0.860 |
| CC | Time constant | $\tau^{\mathrm{FA}}$ | 4 | 8.4665 | -133.5993 | **0.046** | 0.910 | 0.954 | **0.046** |
| CC | Saturation | $S_{\mathrm{FA}}$ | 4 | -0.0134 | 0.6903 | 0.169 | 0.691 | -0.831 | 0.169 |

**Table S7:** Association statistics for post-mortem interval (PMI) versus FA fixation-model parameters in the corpus callosum (CC). Bold $p$-values indicate $p < 0.05$ for the OLS slope.

#### 7.7 Supplementary statistics: PMI associations at discrete tissue conditions

##### 7.7.1 PMI dependence across discrete tissue conditions

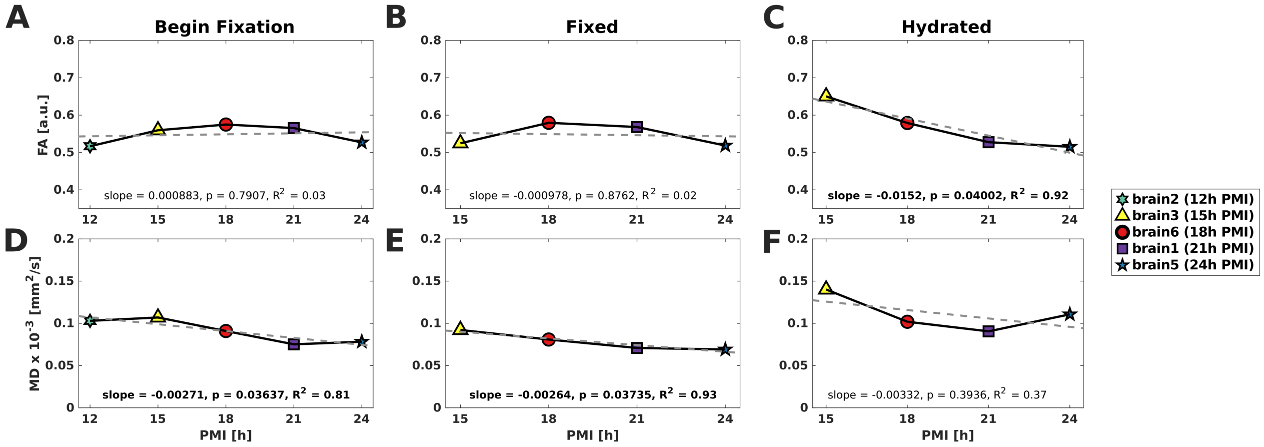


**Figure S10: Post-mortem interval (PMI) in relation to discrete tissue conditions in the corpus callosum (CC):** Brain-wise mean fractional anisotropy (FA) (top row) and mean diffusivity (MD) (bottom row) are plotted against post-mortem interval (PMI) (hours). Columns correspond to beginning of fixation (**A**, **D**; $n = 5$), fixed (**B**, **E**; $n = 4$), and hydrated (**C**, **F**; $n = 4$) tissue. Markers denote individual brains. The gray dashed line in each panel shows the OLS regression. Insets report the slope and corresponding $p$-value; statistically significant results ($p < 0.05$) are shown in bold. Complete OLS and Spearman statistics are provided in Supporting Information Table S8.

At discrete tissue conditions, PMI effects were parameter- and condition-dependent (Figure S10; Supporting Information Table S8). With brain-wise means, MD decreased with increasing PMI at beginning of fixation and in the fixed state ($p < 0.05$), whereas no clear association was observed after hydration. FA decreased with increasing PMI in the hydrated state ($p < 0.05$), but showed no clear association at beginning of fixation or in the fixed state. Within the observed PMI range (12–24 h), these absolute shifts remained small relative to the in-vivo-to-in-situ change for MD and to the broader handling and fixation effects on FA. Thus, condition-level PMI associations were limited, whereas for FA the PMI-related effects remained more apparent in the fitted fixation kinetics (FA($t_0$) and $\tau$).

Figure S10 summarises brain-wise mean FA and MD versus PMI at beginning of fixation, fixed, and hydrated tissue conditions, with OLS regression lines. Table S8 lists the corresponding OLS slopes, intercepts, $p$-values, and $R^2$, together with Spearman rank correlations, for reproducibility. Associations highlighted in the main figure were defined by the two-sided OLS slope $p$-value ($p < 0.05$).

| Condition | Parameter | $n$ | OLS $m$ | OLS $c$ | OLS $p$ | OLS $R^2$ | Spearman $\rho$ | Spearman $p$ |
|---|---|---|---|---|---|---|---|---|
| Begin fixation | FA | 5 | $8.83 \times 10^{-4}$ | 0.533 | 0.791 | 0.027 | 0.300 | 0.683 |
| Begin fixation | MD | 5 | $-2.71 \times 10^{-3}$ | 0.140 | **0.036** | 0.813 | −0.800 | 0.133 |
| Fixed | FA | 4 | $-9.78 \times 10^{-4}$ | 0.567 | 0.876 | 0.015 | −0.400 | 0.750 |
| Fixed | MD | 4 | $-2.64 \times 10^{-3}$ | 0.130 | **0.037** | 0.927 | −1.000 | 0.083 |
| Hydrated | FA | 4 | $-1.52 \times 10^{-2}$ | 0.865 | **0.040** | 0.922 | −1.000 | 0.083 |
| Hydrated | MD | 4 | $-3.32 \times 10^{-3}$ | 0.176 | 0.394 | 0.368 | −0.400 | 0.750 |

**Table S8:** Association statistics for post-mortem interval (PMI) versus brain-wise mean fractional anisotropy (FA) and mean diffusivity (MD) at discrete tissue conditions in the corpus callosum (CC) ($n = 5$ at beginning of fixation; $n = 4$ at the fixed and hydrated conditions). The OLS regression line can be read as $Y = m\,\mathrm{PMI} + c$, where $Y$ is the diffusion parameter, $m$ is the slope, and $c$ is the intercept. Spearman $\rho$ is reported as a complementary rank-based association. Bold $p$-values indicate $p < 0.05$ for the OLS slope.

#### 7.8 Supplementary analysis: fiber orientation dispersion and fixation parameters

The analysis in the main manuscript relates fixation parameters to fiber orientation dispersion ($\kappa$) using Pearson correlation and ordinary least-squares regression across the 15 brain–group points (three brains, five equally sized $\kappa$ quintiles per brain). This analysis provides the two-dimensional summary shown in Figure 6, but it does not explicitly account for the fact that quintiles originate from the same brains. Therefore, we performed a complementary LME analysis with mean group $\kappa$ as fixed effect and brain as a random intercept:

$$Y \sim \kappa + (1|\text{Brain}), \qquad \text{[S6]}$$

where $Y$ denotes the fitted amplitude, time constant $\tau$, or saturation value. The longitudinal fit parameters were obtained from the AICc/ER-selected null, monoexponential, or biexponential saturation model as described in Section 7.5.1.

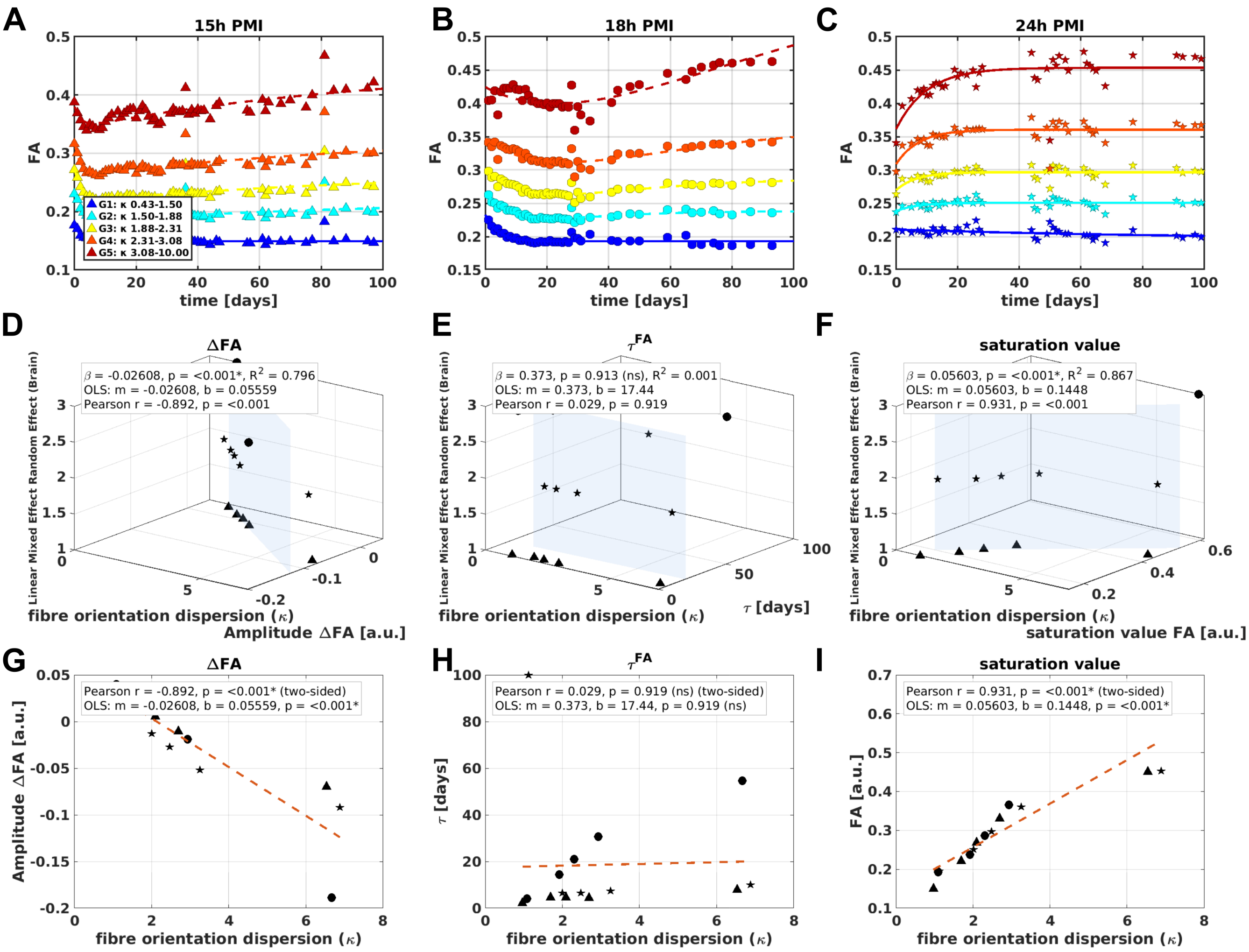


**Figure S11: Supplementary fiber-orientation analysis for fractional anisotropy (FA).** The upper row shows mean FA over fixation time for five equally sized $\kappa$ quintile groups in three brains (15, 18, and 24 h PMI); fitted curves correspond to the AICc/ER-selected saturation model. The middle row shows the complementary LME analysis for fitted amplitude, $\tau$, and saturation FA, with brain included as a random intercept. The bottom row shows the corresponding Pearson correlation and ordinary least-squares regression used for the main-text summary. For FA, the mixed-effects analysis confirms the two-dimensional associations: higher $\kappa$ was associated with a more negative fitted amplitude and higher saturation FA, whereas $\tau$ showed no consistent dependence on $\kappa$.

For FA, the LME analysis supports the same interpretation as the Pearson/OLS analysis in the main text. The fitted amplitude and saturation value showed strong associations with $\kappa$, whereas $\tau$ remained non-significant even after modeling brain as a random intercept. Thus, fiber orientation dispersion primarily affected the magnitude and asymptotic level of the fixation-related FA response, not its time scale.

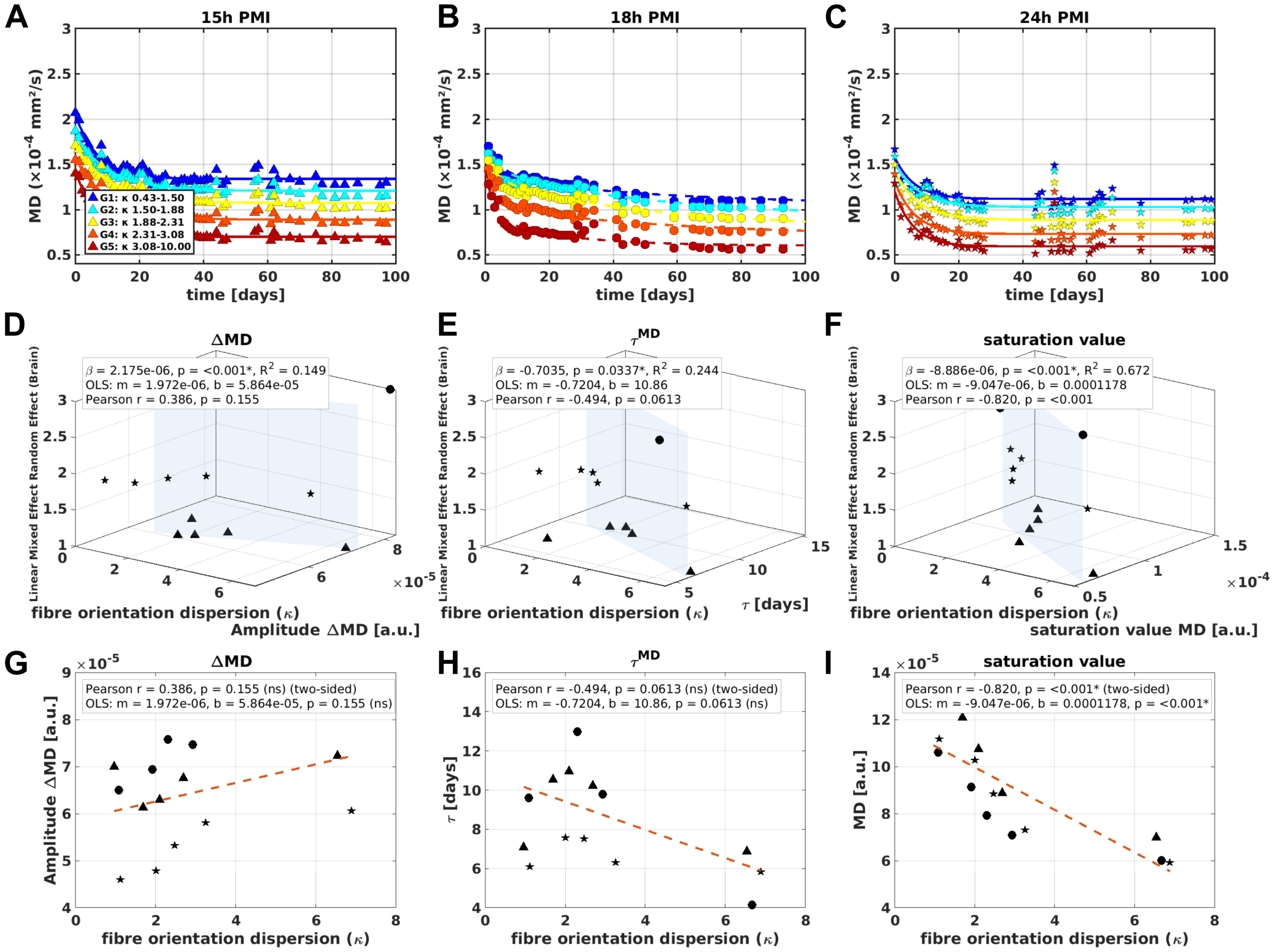


**Figure S12: Supplementary fiber-orientation analysis for mean diffusivity (MD).** The upper row shows mean MD over fixation time for five equally sized $\kappa$ quintile groups in three brains (15, 18, and 24 h PMI); fitted curves correspond to the AICc/ER-selected saturation model. The middle row shows the complementary LME analysis for fitted amplitude, $\tau$, and saturation MD, with brain included as a random intercept. The bottom row shows the corresponding Pearson correlation and ordinary least-squares regression. For MD, the clearest association was observed for the saturation value, which decreased with increasing $\kappa$. The two-dimensional Pearson/OLS analysis showed no significant amplitude association and a trend for $\tau$, whereas the mixed-effects model indicated additional brain-adjusted associations for amplitude and $\tau$.

For MD, only the saturation value was significant in the two-dimensional Pearson/OLS analysis. When brain was included as a random intercept in the LME model, all three fitted parameters were associated with $\kappa$: amplitude, $\tau$, and saturation. The $\tau^{\text{MD}}$ association was the weakest of these effects, while the saturation value remained the most robust finding.

| Parameter | Metric | Pearson $r$ | Pearson $p$ | OLS $m$ | OLS $c$ | OLS $p$ | LME $\beta_\kappa$ | LME $c$ | LME $p$ | LME $R^2_m$ |
|---|---|---|---|---|---|---|---|---|---|---|
| FA amplitude | $\Delta$FA | -0.892 | $7.89 \times 10^{-6}$ | -0.0261 | 0.0556 | $7.89 \times 10^{-6}$ | -0.0261 | 0.0556 | $3.68 \times 10^{-6}$ | 0.796 |
| FA time constant | $\tau^{\text{FA}}$ | 0.029 | 0.919 | 0.373 | 17.442 | 0.919 | 0.373 | 17.442 | 0.913 | 0.001 |
| FA saturation | $S_{\text{FA}}$ | 0.931 | $4.68 \times 10^{-7}$ | 0.0560 | 0.1448 | $4.68 \times 10^{-7}$ | 0.0560 | 0.1448 | $2.06 \times 10^{-7}$ | 0.867 |
| MD amplitude | $\Delta$MD | 0.386 | 0.155 | $1.97 \times 10^{-6}$ | $5.86 \times 10^{-5}$ | 0.155 | $2.18 \times 10^{-6}$ | $5.80 \times 10^{-5}$ | $5.09 \times 10^{-4}$ | 0.149 |
| MD time constant | $\tau^{\text{MD}}$ | -0.494 | 0.061 | -0.720 | 10.856 | 0.061 | -0.703 | 10.806 | 0.034 | 0.244 |
| MD saturation | $S_{\text{MD}}$ | -0.820 | $1.84 \times 10^{-4}$ | $-9.05 \times 10^{-6}$ | $1.18 \times 10^{-4}$ | $1.84 \times 10^{-4}$ | $-8.89 \times 10^{-6}$ | $1.17 \times 10^{-4}$ | $1.24 \times 10^{-5}$ | 0.672 |

**Table S9:** Association statistics for fiber orientation dispersion ($\kappa$) and fixation parameters. Pearson and OLS correspond to the two-dimensional main-text analysis and treat all 15 brain–group points as individual observations. The plotted OLS regression line can be read as $Y = m\kappa + c$, where $Y$ is the fitted fixation parameter, $m$ is the slope, and $c$ is the intercept at $\kappa = 0$. The LME fixed-effect relation is analogous, $Y = \beta_\kappa \kappa + c$, with brain included as a random intercept.
